# A Microneedle Device for Rapid Dermal Interstitial Fluid Sampling

**DOI:** 10.1101/2025.03.13.641882

**Authors:** Andy H. Hung, Netra U. Rajesh, Abel Bermudez, Stephanie M. Boczek, Fernando J. Garcia-Marqués, Yee Lin Tan, Jihyun (Luna) Hwang, Prima Dewi Sinawang, Dan Ilyin, Gunilla B. Jacobson, Utkan Demirci, Steven P. Poplack, Sharon J. Pitteri, Joseph M. DeSimone

## Abstract

Dermal interstitial fluid (ISF) offers a promising alternative to invasive blood tests and opportunities for novel skin diagnostics. Progress in both the understanding and adoption of ISF tests is hindered by sampling challenges, including lengthy collection times, non-negligible failure rates, variable collection volumes, and inconsistent bioanalyte levels. The causes of many of these issues are not well understood. We demonstrate a microneedle device that is several times faster than state-of-the-art, collecting an average of 15.5 mg of ISF in 5 minutes in humans with near-zero failure rate. This improvement was achieved by designing the spatial pressure gradient driving ISF flow. The influence of penetration depth, collection time, pressure, and age on ISF collection was elucidated, with Darcy’s law explaining multiple observations. A data-driven acceptance criterion of <1% blood contamination for ISF is proposed. The device and findings presented will empower researchers to better conduct robust studies in the development of ISF diagnostics.

## Introduction

Dermal interstitial fluid (ISF) occupies the space between cells in the skin and exchanges biomolecules with blood, cells, and lymph^1,2^. Nearly all plasma analytes are present in ISF, with a small fraction of analytes exclusive to ISF, presumably from the skin microenvironment^3–7^. These biomarkers position ISF as a promising minimally invasive diagnostic alternative to blood and a tool for evaluating skin conditions. However, the development of ISF testing is hindered by the absence of a sufficiently fast and robust sampling method, and poorly understood variabilities in the collection volumes and measured biomarker levels^3,4,8,9^.

The main challenges in ISF sampling stem from its residence in the spaces within the skin extracellular matrix (ECM). First, more than 40% of ISF is noncovalently bound water that cannot be sampled, with an estimated accessible ISF of only 120 μL per cm^2^ of skin in the thickest regions of the dermis^3^. In practice, current ISF sampling devices collect 1-20 μL^4,6,8,10^. Second, the ECM creates high resistance to ISF flow during collection^2^, leading to lengthy procedures of 20 minutes or more^4,6,8,10^. Third, the ECM is hypothesized to impose a chromatographic effect on ISF analytes during flow, which would cause the large and small analytes to separate, resulting in inconsistencies in their measured levels^3^. Finally, the ECM changes depending on individual age, gender, ethnicity, body region, and lifestyle, further adding to the variability in ISF sampling. These complexities make ISF sampling more challenging than blood sampling.

Traditional ISF sampling methods are invasive, including wick extraction^11^, suction blister formation^12^, and microdialysis^13^. Microneedle-based methods are minimally invasive, but historically suffered from low collection volumes, typically <1 μL^14^. More recently, a key insight to couple pressure with microneedles dramatically improved the collection speed and volume of these devices^6,14,15^, catapulting them to the foremost current approach in ISF research. Common approaches include puncturing skin with a microneedle array followed by vacuum suction^8,14,15^ and applying positive pressure around hollow microneedles to drive ISF collection^6,10^. In the latest iteration, Jiang et al. achieved an impressive mean collection volume of 20.8 µL in 25 min following the vacuum approach utilizing 1200-1600 puncture sites^8^.

Building on these pioneering works, we designed a 3D-printed^16–18^ microneedle ISF sampling device that collects a mean of 15.5 mg in 5 min utilizing 154 puncture sites in humans. This performance was achieved by engineering from first principles following Darcy’s Law, which describes fluid flow through a porous medium (Fig. 1a)^2,14,19^. Specifically, we focused on designing the *distribution* of pressure gradients rather than the magnitude of applied pressure. The concept (Fig. 1b) is to puncture the skin using a microneedle array patch (MAP) through a collection plate with aligned perforations. The MAP is removed, and the plate is pushed down, applying pressure in *all* regions where flow should be directed away, and zero pressure at the puncture sites where flow should be directed to. This approach optimizes the spatial pressure gradient in the skin to maximize ISF flow. ISF flowing into the puncture sites are collected by surface tension in the plate perforations. We call this device the Puncture-Out-and-Press device, or POP.

**Figure 1:**
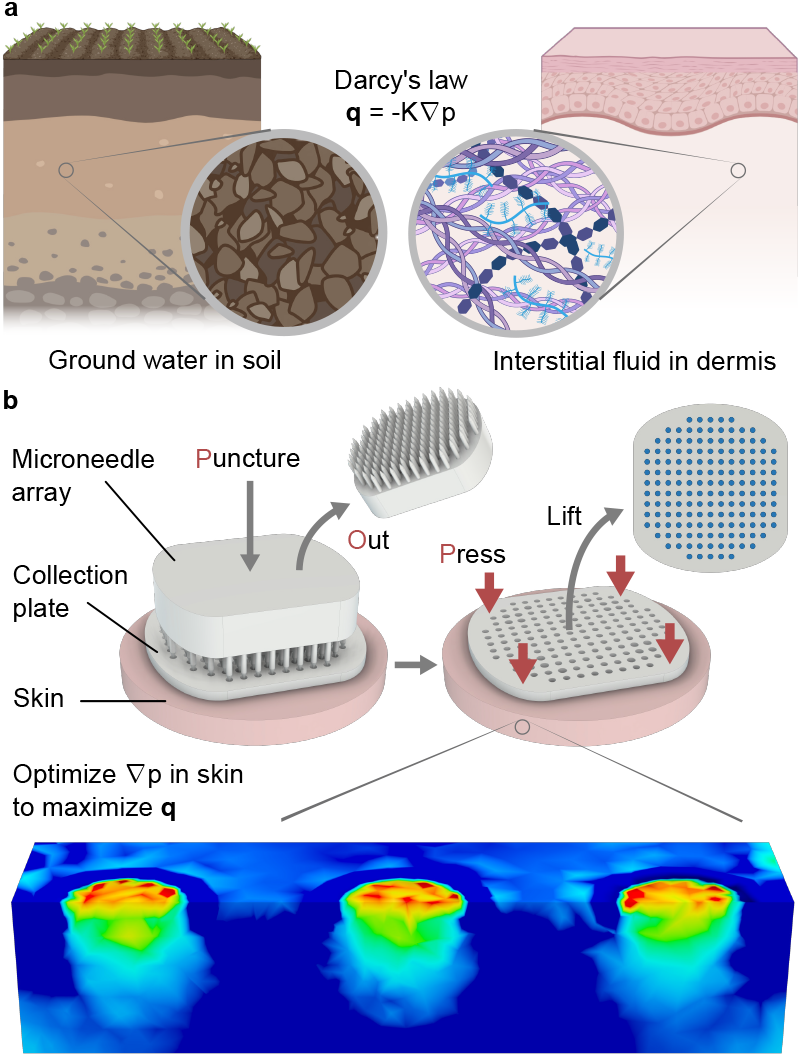
Designing by Darcy’s Law. **a** Derived from groundwater flow in soil, Darcy’s law describes flow in a porous medium, which applies to ISF flow in dermis. **q** is flux vector, K is conductivity, ∇p is pressure gradient. **b** Puncture-Out-and-Press (POP) device concept. The key innovation is in designing the distribution of applied pressure using a perforated collection plate. The optimized ∇p within the dermis drives ISF flow to the puncture sites, which is collected in the plate by surface tension.

Leveraging the larger collection volumes and near-zero collection failure rate of POP, we conducted studies to increase the mechanistic understanding of ISF sampling, quantify the effect of blood contamination in ISF, and revisit the long-held notion that protein transport into ISF is size-dependent^3,20^. These studies provide further foundation for improving the robustness and predictability of ISF sampling and bring the field closer to widespread use of ISF tests.

## Results

### POP Design Dimensions

The base POP design featured a MAP with 7 × 5 solid microneedles with 400 µm diameter conical tips, and a collection plate with 500 µm perforations. When mated, 1100 µm of the microneedles are exposed. Depending on experimental needs, slightly different versions of POP were 3D-printed (Fig. S1, S2, S3).

### Spatial Pressure Gradient Design

To test the idea that spatial pressure gradient modulates ISF flow, we altered pressure distributions in *ex vivo* human skin by varying the use configuration and design details of POP, then compared ISF collection. Specifically, we tested the POP design, a needle-in variant that applied pressure on the MAP and plate together without MAP removal, a vacuum variant, and a variant that applied pressure in a ring distribution (Fig. 2). Because the needle-in variant required a hollow MAP to collect ISF, hollow MAPs were used in all variants to eliminate skin puncture as a confounding variable. POP performed similarly regardless of whether a solid or hollow MAP was used for skin puncturing (Fig. S4). Finite element analysis (FEA) employing a simplified dermis model illustrated the different pressure gradients.

**Figure 2:**
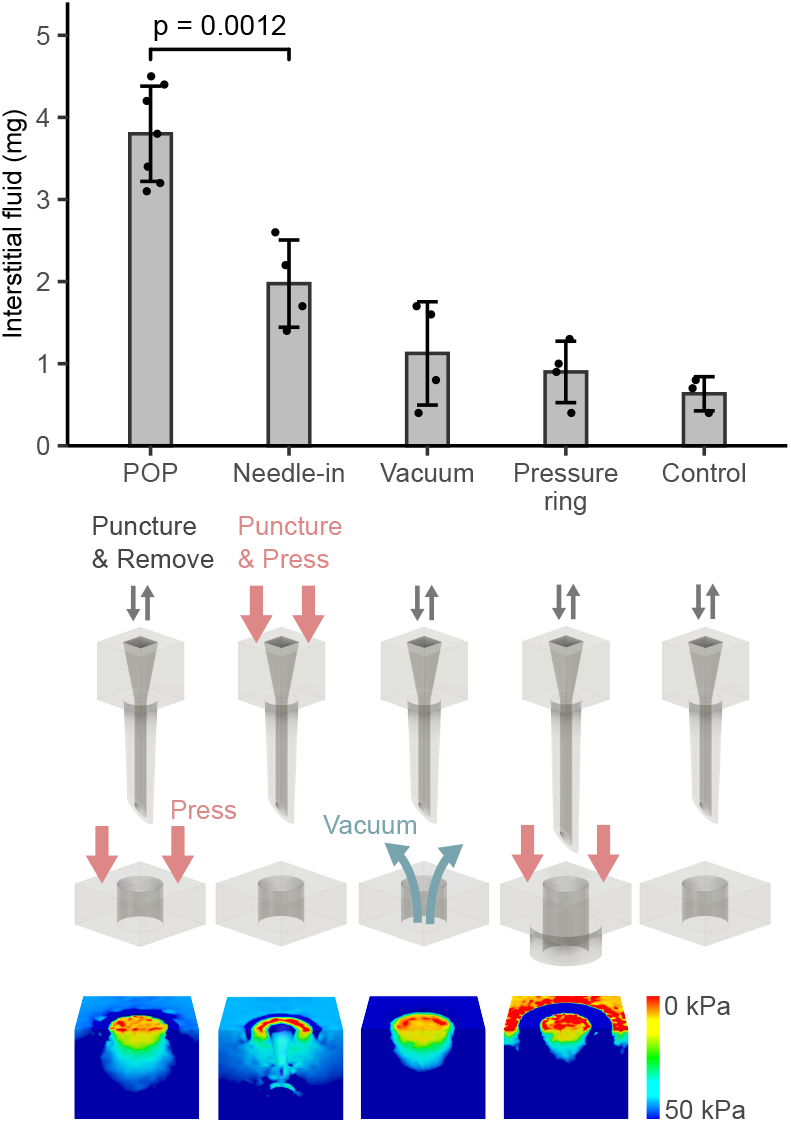
Spatial pressure gradient modulates ISF flow. ISF collected from *ex vivo* human skin using POP outperformed the needle-in variant, the vacuum variant, and the pressure ring variant. A control without pressure application was included. FEA simulations illustrate the tissue pressure distribution for each design, explaining ISF flow variations. Color scale shows relative pressure. Flow moves from cool to warm colors. Pressure was applied with a 500 g weight, equivalent to 50 kPa in POP. The vacuum applied was -64 kPa. Collection time was 5 minutes. Statistical analysis used a student’s t-test.

POP extracted 1.9 to 4 times more ISF at 50 kPa compared to alternatives. The simulated pressure distribution for POP showed the entire tissue to be in compression except at the puncture site, as designed.

In the needle-in variant, the MAP remained in place, and pressure was applied on the MAP. ISF flowed into the hollow microneedle instead of the puncture site. Compared to POP, ISF flow was reduced because tissue distension by the microneedle residing in the puncture site diminished the pressure gradient between the puncture site and surrounding tissue.

In the vacuum variant, -64 kPa was applied to the plate after MAP removal. While FEA predicted a greater pressure gradient than POP, experimental results did not match the prediction, likely due to difficulties in maintaining vacuum seals across 7 × 5 openings. To investigate this discrepancy, we applied vacuum directly on skin following MAP puncture. This ensured a reliable seal but eliminated pressure gradients in the lateral dimensions. In this setup, the collection rate sometimes matched POP (Fig. S5).

In the pressure ring variant, ring protrusions on the collection plate distributed pressure in a ring pattern. The MAP was lengthened to account for the additional height from the ring protrusions (Fig. S3). Compared to POP, collection efficiency was reduced because ISF flowed both towards and away from the puncture site, as shown by the pressure distribution from FEA.

Overall, this series of comparisons underscores the primary role of the designed spatial pressure gradient in achieving POP’s rapid ISF collection rate.

### Mechanistic Investigations

Substantial variabilities observed in the collection volume, fluid content, and collection failure rate of ISF sampling devices pose challenges for ISF diagnostics^4,6,8^. Gaps remain in the mechanistic understanding of ISF sampling, hindering the control of variability^9^. To this end, we investigated the impact of penetration depth, collection time, and pressure on the quantity and quality of collected ISF. Previous studies have focused on empirical optimization^6,8,14^ and provided limited mechanistic insights due to biological heterogeneity or small collection volumes. Here, we used *ex vivo* skin from a single donor for each experiment to minimize the confounding effects of tissue-to-tissue variation; combined with the increased experimental precision from larger ISF volumes, instructive patterns in the data were identified.

### Penetration Depth

To understand the role of penetration depth in ISF collection, we compared POP on porcine ear and back skin (Fig. 3a, S6, S7). The mean penetration depth was 480 μm on the softer ear skin and 169 μm on the tougher back skin. Histological examination showed that punctures reached regions with larger interstitial spaces in the ear skin, but not in the back. This difference led to more effective ISF collection from the pig ear skin. A control experiment confirmed similar hydration and conductivity in the ISF-rich regions of both skin types (Fig. S8). These results provide strong empirical support that sufficient penetration depth into the ISF-rich region of the skin is necessary for efficient ISF collection.

**Figure 3:**
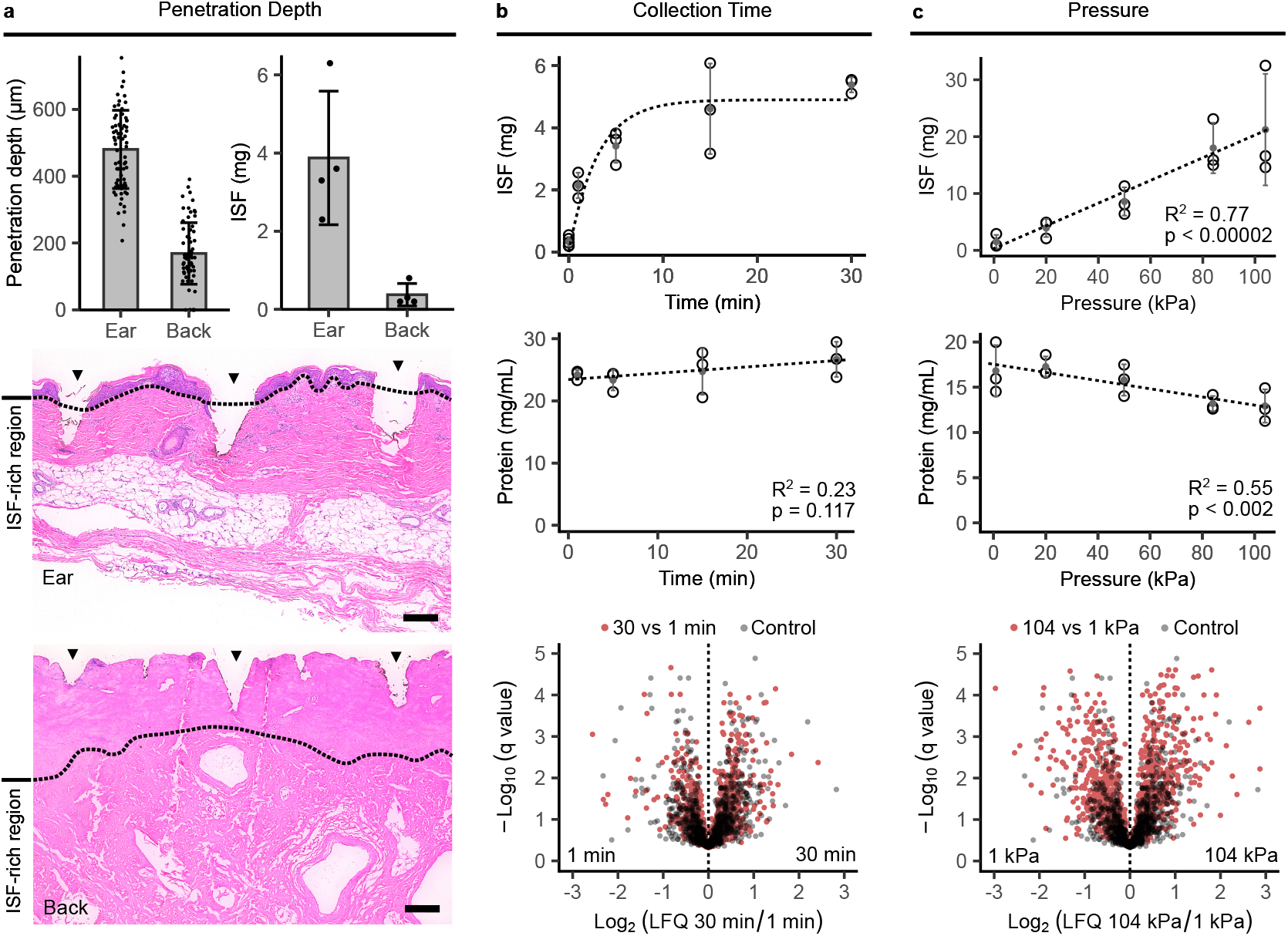
ISF quantity and content change with collection parameters. **a** Comparison of POP on porcine ear and back skin demonstrates that efficient ISF collection requires sufficient penetration into the ISF-rich regions of skin. Arrowheads show puncture sites. Scale bars = 200 µm. **b** ISF flow decreased over time due to reduced conductivity from tissue compaction. No link was evident between collection time and ISF proteome. Pressure was 50 kPa. Controls were collected without pressure. **c** ISF flow increased linearly with pressure following Darcy’s law. Pressure reduced protein concentration and increased dispersion in volcano plot analysis, indicating separation of protein species by the ECM. Controls were the same as in **b**. Statistical analysis used a non-linear model based on Darcy’s law, linear regression, and the Storey procedure.

The implication is that to enhance sampling consistency, a device should be designed to ensure that most punctures reach the ISF-rich region of the skin. The human epidermis ranges 50-250 μm in thickness^21^, and penetration can vary up to 200-300 μm on a single patch (Fig. 3a). Designing for the average case without accounting for these distributions can lead to many punctures not reaching effective depths. Inadequate penetration may explain some collection volume variability and instances of collection failure in the literature.

### Collection Time

Next, we examined collection time using POP on *ex vivo* human skin, maintaining a constant pressure of 50 kPa (Fig. 3b). Intuitively, as collection proceeds and the solid tissue fraction increases, flow would slow due to reduced conductivity. To a first order approximation, conductivity is modeled as a linear function of the ISF volume fraction^2,22,23^, resulting in:

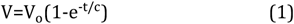

where V is the volume of ISF extracted, V_0_ is the maximum possible volume extraction under the specific collection conditions, t is collection time, and c is a constant. This simplified model effectively approximates ISF collection over time, reaffirming Darcy’s law as a valuable heuristic in understanding ISF collection.

In addition to assessing ISF quantity, we evaluated its quality. A concern with pressure-based ISF collection is that the ECM might restrict large molecule extraction, affecting the accuracy of ISF tests^3^. This hypothesis, rooted in suction blister fluid data^24^, has not been explored with microneedle devices. The hypothesis predicts lower protein concentration and increased abundance of low molecular weight species in ISF over longer collection times. However, no such correlation was found between collection time and ISF protein concentration (Fig. 3b). Similarly, volcano plot analysis showed that ISF proteomes from the 30-minute and 1-minute collections were similar to controls collected without pressure application. Thus, varying collection times provided no evidence that the ECM acts as a size-based filter for proteins.

### Pressure

We conducted similar analyses with pressure using POP on *ex vivo* human skin, keeping the collection time constant at 5 minutes (Fig. 3c). A strong positive linear correlation emerged between applied pressure and ISF collection, aligning with Darcy’s law and previous studies^14^. Although a sublinear relationship was anticipated based on decreased hydraulic conductivity at larger collection volumes, this was not convincingly observed. One possibility is that abundant skin hydration in the specimen allowed conductivity to be maintained until more than 15 mg of fluid was extracted, as the 104 kPa data point appeared to plateau.

Unlike the time series findings, ISF protein concentration decreased with increasing pressure, consistent with the ECM filtration theory^3^. The change was small, decreasing by 23% from 17 mg/mL to 13 mg/mL as pressure increased from 0.9 kPa to 104 kPa. Volcano plot analysis confirmed this finding, showing that 21% of proteins significantly differed between ISF collected at 104 kPa and 0.9 kPa at a false discovery rate (FDR) of 0.01. In contrast, only 8% of proteins significantly differed in the control. Analysis by molecular weight showed a subtle enrichment for high molecular weight species under high pressure (Fig. S9). This supports a size exclusion model, where larger molecules move faster through the matrix due to shorter path lengths in the large pores, rather than a filtration model where large molecules are trapped. This phenomenon can contribute to variability in detected ISF composition.

### Human Pilot Study

POP was evaluated in a pilot human study with 10 subjects (Table S1), with safety and pain as the primary outcomes and collection volume and proteome measurement as the secondary outcomes. 5 participants underwent finger prick blood collection on the same arm immediately before ISF collection, with an average volume of 4.2 ± 1.0 µL. Sham collections without MAP puncture were performed on three additional subjects to ensure that any downstream analysis was reflective of ISF and not skin surface contaminants (Table S2).

The clinical study version of POP used a MAP with 154 solid microneedles, a topper to ensure uniform pressure on the collection plate after MAP removal, and a stand for positioning the spring-loaded applicator at a controlled height (Fig. 4a, b, S1, S10). Prior to study enrollment, the device was shown to undergo deformation without fracture during mechanical failure (Fig. S11), and monomer leaching was validated to be absent (Fig. S12). After puncture and replacement of MAP by the topper, a force sensor was used to maintain the applied pressure at 750 g, equal to 25 kPa. This pressure was selected in view of normal systolic blood pressure (16 kPa) to reduce risks of ischemic skin damage and pressure ulcers. For comparison, vacuum-based devices typically use -50 kPa^4,8^.

**Figure 4:**
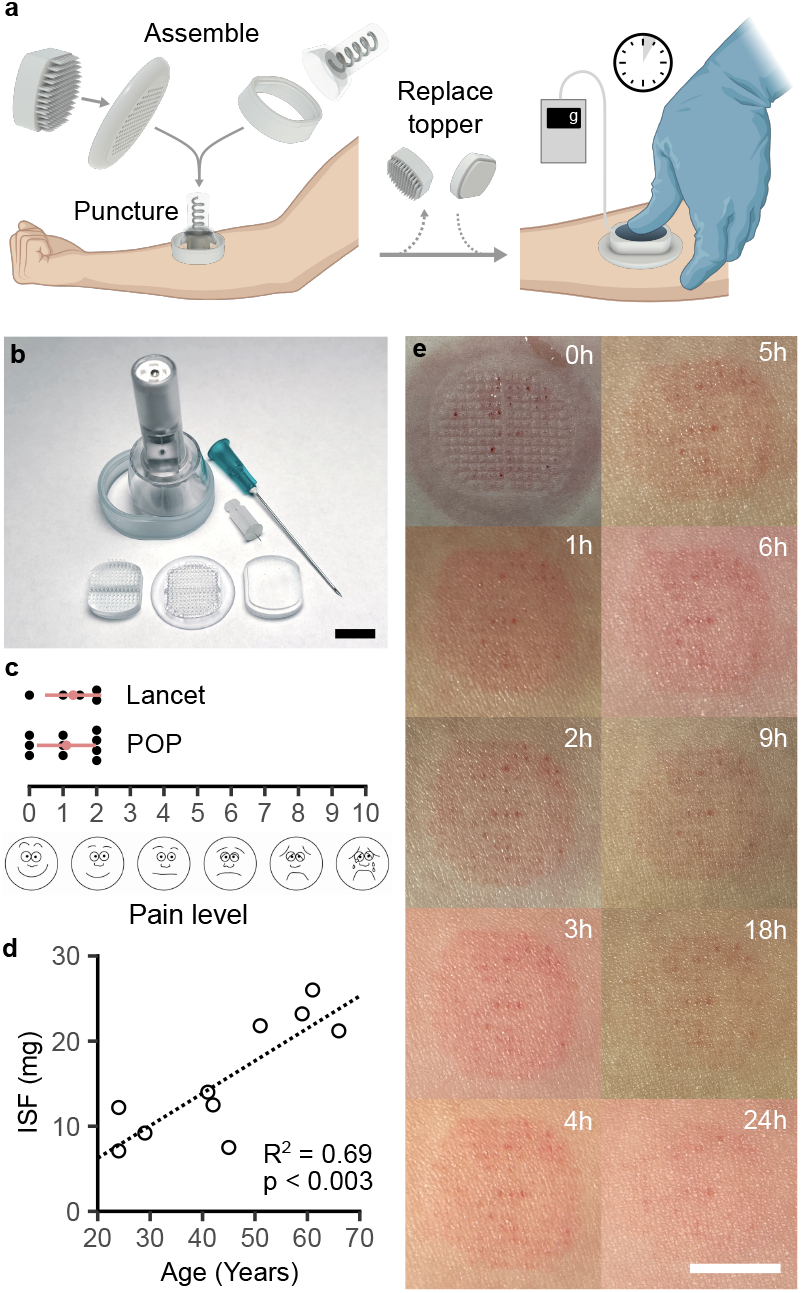
Pilot human study. **a** ISF collection using POP in humans. **b** Components of the POP device, in comparison to a 23G needle and a 33G lancet. **c** Pain levels on the visual analog scale from POP were comparable to a finger prick. **d** Age explains 69% of the variance in ISF collection amounts. **e** POP puncture site evolution over 24 hours. Statistical analysis used linear regression. Scale bars = 10 mm.

On average, 15.5 ± 7 mg of ISF was collected in 5 minutes, with a pain score of 1.1 ± 0.9 on the visual analogue scale. Pain from POP was similar to a finger prick (Fig. 4c). Immediately post-collection, trace blood and a pressure imprint were visible. Erythema nearly resolved within 24 hours (Fig. 4e, S13). No other adverse events occurred. A strong positive correlation between age and ISF collection amount was observed, with age accounting for 69% of the variance (Fig. 4d). Based on Darcy’s law, this could be due to an increased hydraulic conductivity from ECM loss with age. To explore this idea, increased ISF collection was shown in *ex vivo* human skin pre-treated with hyaluronidase to degrade part of the ECM (Fig. S14). Thus, age is a newly uncovered demographic factor that can explain ISF collection variability.

### Blood Contamination

Understanding blood contamination is as crucial for ISF tests as understanding hemolytic contamination is for blood tests^25^. Despite the common occurrence of blood contamination in ISF sampling^10,14^, there has been no study to date analyzing its effects. While device engineering can mitigate contamination, complete elimination is unlikely as adoption of ISF testing increases. Therefore, data is needed to establish acceptance criteria for ISF samples. We present the first quantitative study of errors resulting from blood contamination in ISF at the proteomic level, and examine centrifugation and statistical removal of red blood cell (RBC) proteins as remediation strategies (Fig. 5a).

**Figure 5:**
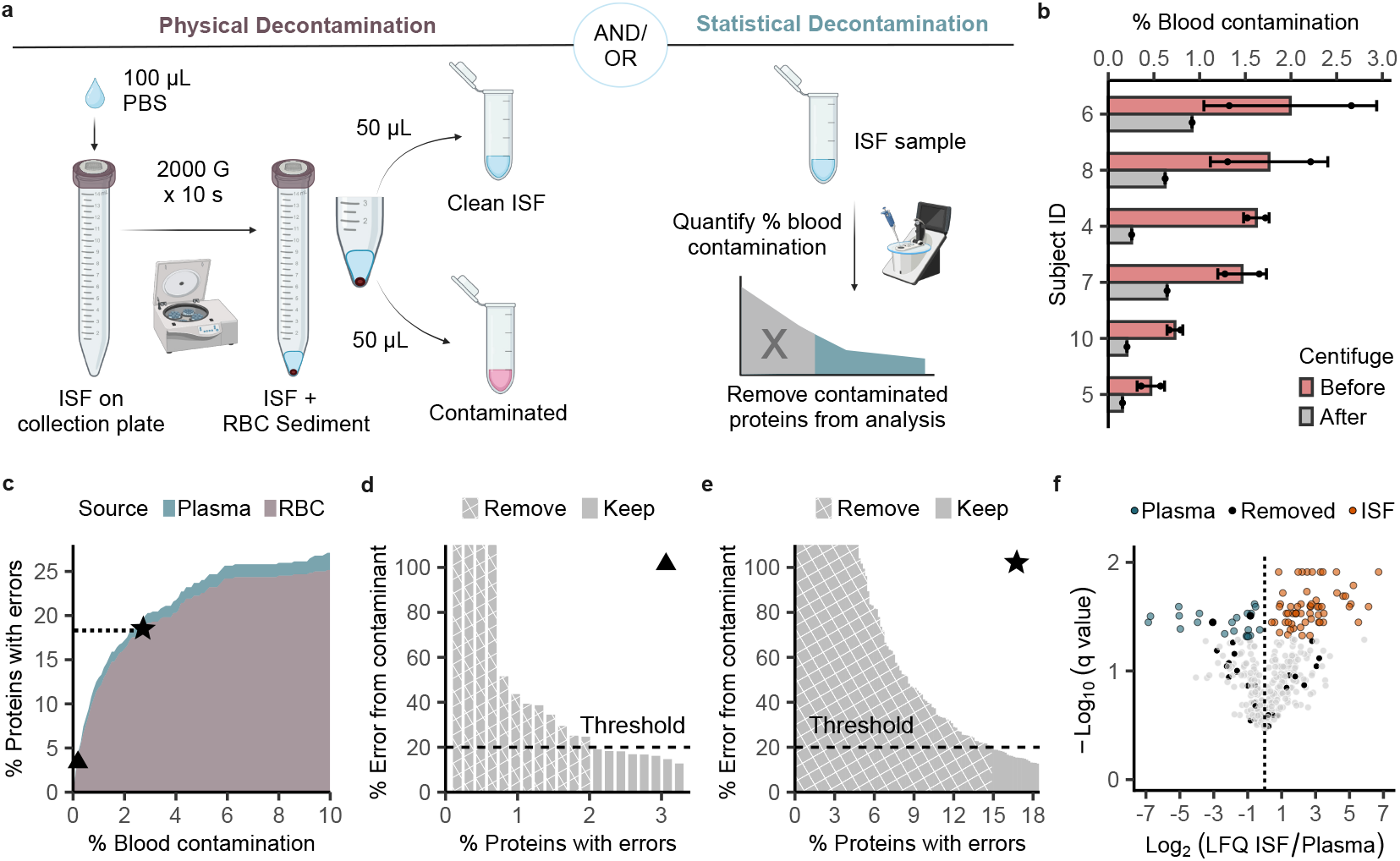
Blood contamination in ISF. **a** Contamination can be physically mitigated by collecting only the supernatant after centrifugation of ISF from the collection plate. It can be statistically mitigated by excluding contaminated proteins from data analysis. **b** Blood contamination was <2% by volume after POP collection and <1% after centrifugation. Residual contamination was due to hemolysis. **c** Quantitative relationship between the volume percentage of blood contamination and the percentage of the proteome contaminated. Most contaminants originate from RBCs. A protein is defined as contaminated if its quantitation has >10% error due to a contaminant. **d** At 0.16% blood contamination, 20 proteins representing 3.2% of the measured proteome are contaminated, with 4 proteins showing >100% error. Either all proteins can be excluded from data analysis, or a less stringent threshold can be set based on the analysis goal. **e** At 2.7% blood contamination, >18% of the proteome is contaminated. **f** Applying the statistical decontamination pipeline to volcano plot analysis comparing ISF to plasma using the most stringent threshold, only 2 out of 81 statistically significant proteins are excluded from analysis. An acceptance criterion of <1% for ISF blood contamination appears reasonable. Statistical analysis used the Storey procedure.

ISF collected by POP showed blood contamination below 2% by volume. During processing, brief centrifugation simultaneously extracted ISF from the collection plate and separated ISF from RBCs, reducing contamination to below 1% (Fig. 5b). The residual contamination originated from hemolysis during collection and handling based on the absence of RBCs on microscopy.

To elucidate the relationship between blood contamination and proteome quantification error, we performed relative quantification of proteins in plasma, ISF, and RBCs.

Errors caused by contaminants can be derived from a linear mix model of ISF and contaminant levels:

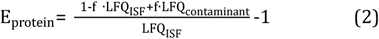

E_protein_ is the percentage error for a particular protein, f is the percentage blood contamination by volume, and LFQ is the label-free quantification derived from mass spectrometry, where the contaminant can be either plasma or RBC. When the error introduced by contaminants exceeds measurement error, conservatively estimated at 10% (Fig. S15), a protein is defined as contaminated.

Results show the non-linear correlation between blood contamination and the affected proteome fraction, with RBCs as the main contamination source due to plasma’s compositional similarity with ISF (Fig. 5c). In our study, the contribution from plasma was ignored because no contamination exceeded 2%.

At a more detailed level, individual protein errors at different blood contamination levels can be estimated. At 0.16% contamination, 4 proteins representing 0.6% of the quantified proteome showed errors >100% (Fig. 5d); At 2.7% contamination, 4.8% of the quantified proteome showed errors >100% (Fig. 5e). This data provides a comprehensive estimation of errors due to blood contamination.

To remove contaminant RBC proteins from downstream analysis, hemoglobin levels in ISF are measured to estimate blood contamination, followed by the removal of all proteins identified as contaminated (Fig. S16, S17). Alternatively, a flexible error threshold can be applied for analysis comparing ISF and plasma, blood contamination showed minimal impact, with only 2 out of 81 significant proteins removed (Fig. 5f). Taken together, an acceptance criterion of <1% blood contamination for ISF appears reasonable, and is close to the acceptance criterion of 0.3 - 2.0% for hemolytic contamination in blood tests, depending on the desired measurement^25^.

### Analytical Quality of ISF Collected by POP

To evaluate the analytical quality of ISF collected by POP, we performed proteomic analysis by mass spectrometry. The mean ISF protein concentration was 29 ± 7 mg/mL, or 44% of the mean plasma protein concentration (Fig. 6c), consistent with the previously reported range of 20-60 mg/mL^8,11,26^. The top 20 proteins substantially overlap in ISF and plasma and represent >85% of the proteome by iBAQ^27^ in both (Fig. 6a). This result suggests that the high-abundance species would mask the low-abundance species equally in ISF and plasma during proteome measurement, contrary to previous conjectures^28^. By relative quantification, more protein species were over-represented in the ISF proteome compared to plasma, similar to reports by others (Fig. 5f)^5,8^. Albumin constitutes a larger fraction of the ISF proteome compared to plasma, with apolipoproteins being less prevalent in ISF (Fig. 6a,b). This result is consistent with the ultrafiltration model of plasma-to-ISF transport^3,24,29,30^, with reduced transport of large lipoproteins (Fig. S18) into ISF compared to the relatively smaller albumin. For immunoglobulins, the difference between plasma and ISF is likely overestimated (Fig. 6a) due to the significant variance in plasma fractions between individuals (Fig. S19). Relative quantification reveals that dominant immunoglobulin chains, such as IGHG1, IGHG2, and IGK/IGKC, exhibit similar proteomic abundance in both fluids, as previously reported (Fig. 6b, S19, Table S3)^31^. Finally, 149 of the identified ISF proteins are in the National Cancer Institute Early Detection Research Network Biomarker database, similar to findings by Jiang, et al^8^. A majority of these proteins are potential breast and ovarian cancer biomarkers (Fig. S20). Overall, these findings are consistent with previously published data, validating the utility of ISF collected by POP.

**Figure 6:**
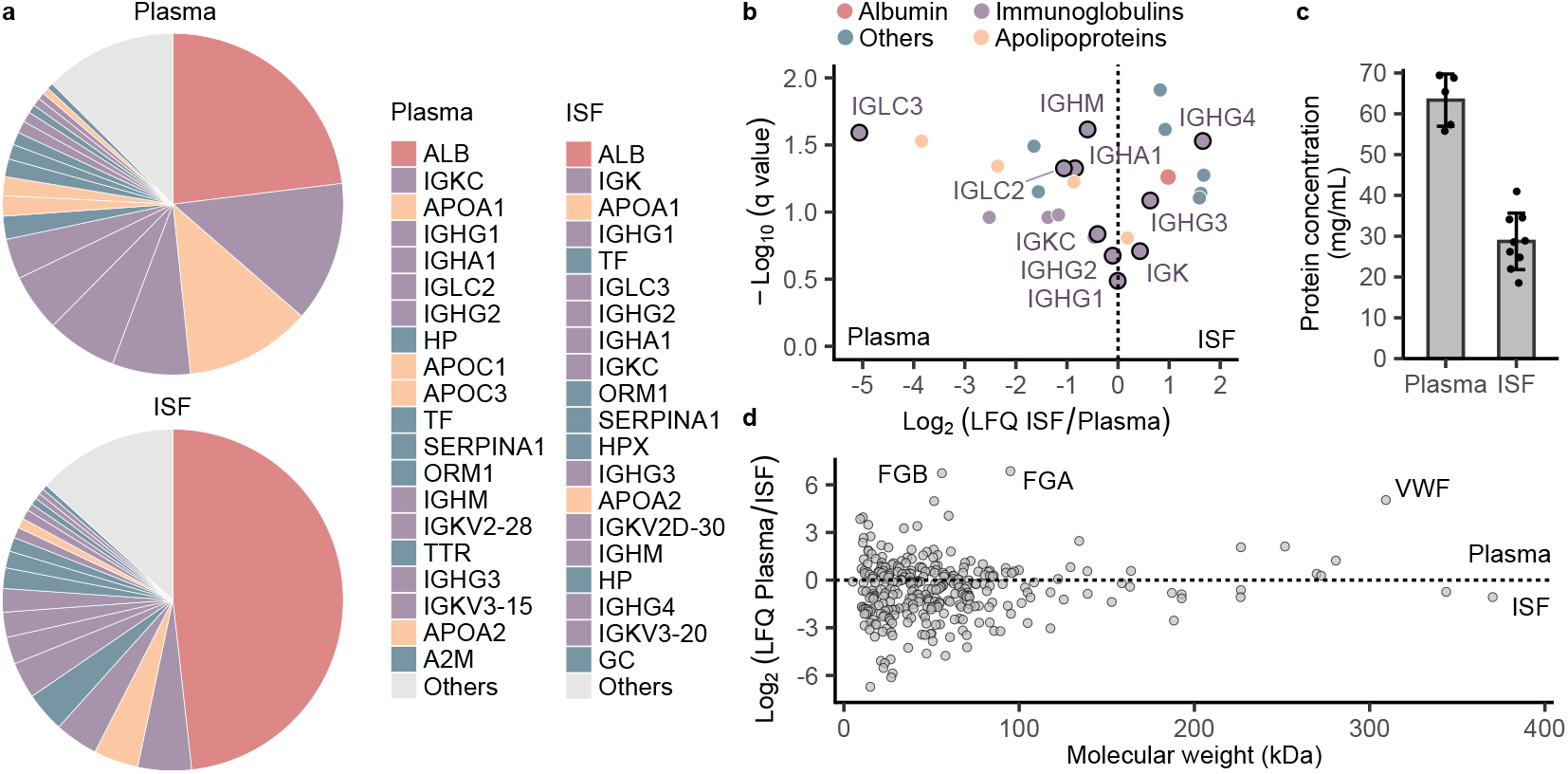
ISF protein composition relative to plasma by mass spectrometry. **a** Comparison of the most abundant proteins in human plasma and ISF collected with POP, by iBAQ (N = 5). Albumin is over-represented in the ISF proteome relative to plasma. Apolipoproteins are under-represented. Immunoglobulin differences between ISF and plasma are likely overestimated. **b** Relative quantification confirms the findings from iBAQ analysis for albumin and apolipoproteins. The trend is less clear for immunoglobulins, with no statistically significant difference between ISF and plasma for major classes such as IGHG1, IGHG2, and IGK/IGKC. **c** Protein concentration is lower in ISF compared to plasma. **d** Plasma-ISF partitioning is not dependent on molecular weight. The most over-represented plasma proteins are coagulation proteins. Data does not support the ultrafiltration model. Statistical analysis used the Storey procedure.

### Revisiting the Ultrafiltration Model

A novel finding in our ISF analysis is the absence of a clear relationship between plasma-ISF partitioning and molecular weight (Fig. 6d). The plasma proteome does not show enrichment for high molecular weight proteins compared to ISF; instead, the most over-represented plasma proteins are coagulation proteins. This finding contradicts the ultrafiltration model of molecular transport into ISF, which posits that small molecules freely pass through the endothelial barrier from plasma into ISF, while large molecules are increasingly restricted^3,24,29,30^. Although this model accurately describes lipoprotein particle partitioning between plasma and ISF (Fig. 6b), its predictive power is poor at the proteome level. For example, immunoglobulins, known to be some of the largest proteins, are not decisively enriched in plasma relative to ISF (Fig. 6b). Proteins dynamically fold and unfold, raising questions about their adherence to transport laws derived in part using dextran^32^ and lipoprotein data^29,30^. The notion that molecular weight dictates endothelial transport originated from the observation that the ratio of suction blister fluid-to-plasma analyte concentration decreases as molecular weight increases^29^. However, this type of study overlooks the fact that up to two-thirds of ISF in skin is inaccessible to macromolecules due to steric exclusion by hyaluronate and collagen^2,33^. In plasma protein concentration measurements, the cellular fraction inaccessible to proteins is routinely excluded. A similar exclusion of the inaccessible ISF fraction is necessary for a valid comparison when measuring ISF proteins, but this is rarely done because the inaccessible fraction is not obvious after collection. Consequently, high molecular weight protein levels in ISF appear artificially low compared to plasma (Fig. S21). Thus, molecular weight may play a less dominant role than previously thought in protein plasma-ISF partitioning relative to factors like charge, glycosylation, and hydrophobicity. Given the frequent invocation of molecular weight in ISF study interpretation, revisiting the application of the ultrafiltration model to proteins with more comprehensive studies would be beneficial.

## Discussion

In this work, we introduced POP, a minimally invasive device for rapid and robust dermal ISF sampling capable of collecting >10 mg in 5 min in humans. Darcy’s law, which has been underutilized in the understanding of ISF sampling, was found to be powerful both in the design of devices and the explanation of observations. We established empirical evidence that penetration depth, pressure, and age at least partially account for some of the puzzling variabilities seen in ISF collection volumes and bioanalyte levels. Much of the variabilities relate back to the effect of the ECM on flow and bioanalyte separation. An analysis of the errors caused by blood contamination supports <1% contamination as an acceptance criterion for ISF, with two suggested mitigation strategies.

In the future, POP enhancements may include a minicentrifuge adapter for clinical portability, a porous collection plate layer for enhanced ISF retention, and a monolithic design with a latticed MAP^18^ for improved usability (Fig. 7, S22, S23). The new device in this report, together with the findings uncovering previously unknown or unconfirmed factors contributing to ISF variability, lays the foundation for greater scale and robustness in future ISF clinical research.

**Figure 7:**
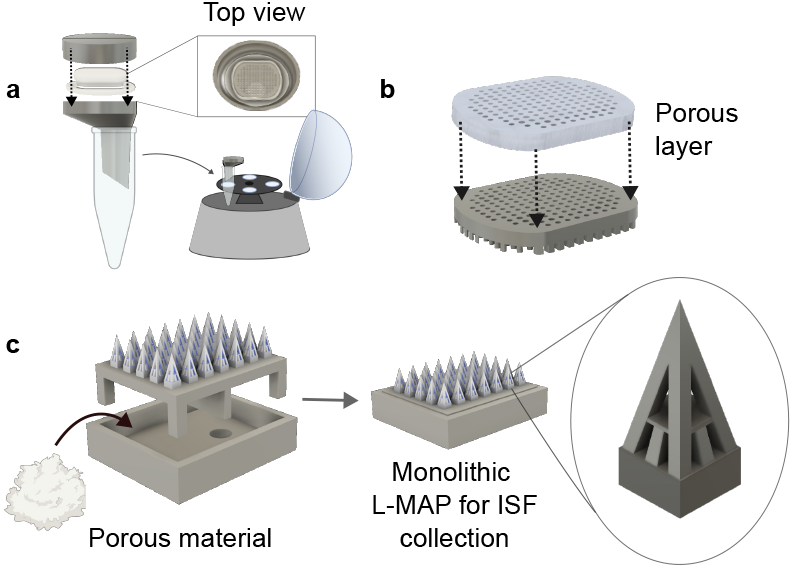
Future Directions. **a** Collection plate adapter for ISF centrifugation into a microcentrifuge tube enables clinical portability. **b** Addition of a porous layer on the collection plate to improve ISF retention. **c** Monolithic latticed MAP design simplifies the sampling procedure.

## Acknowledgements

This work was funded by a Stanford Precision Health and Integrated Diagnostics Center Pilot Grant, the Stanford Cancer Imaging Training Grant T32CA00969531, James D. Plummer Graduate Fellowship, EDGE Doctoral Fellowship Program, Summer First Fellowship Program, Dean’s Office of the Stanford School of Engineering, Cancer Imaging & Early Detection Award, Canary-ACED Graduate Fellowship, and Stanford Bio-X Interdisciplinary Initiatives Program Seed Grant. We thank the Stanford Department of Radiology Office of Clinical Trials and Translational Research for support, and Essy Mathew, Ian Coates, Madison Driskill, Jean Kwak, Jason Kronenfeld, John Tumbleston, Max Saccone, Maria Dulay, Roger Wise, and Professor Mark Prausnitz for helpful discussions.

## Author Contributions

A.H.H. conceptualized the work, acquired, analyzed, and interpreted the data, created software, and wrote the paper. N.U.R. conceptualized the work, acquired, analyzed, and interpreted the data, and wrote the paper. A.B. acquired and analyzed the data. S.M.B. acquired, analyzed, and interpreted the data. F.J.G. analyzed and interpreted the data and created software. Y.L.T. conceptualized the work and acquired, analyzed, and interpreted the data. J.L.H. acquired and analyzed the data. P.D.S acquired and analyzed the data. D.I. conceptualized the work and interpreted the data. G.B.J. conceptualized the work and interpreted the data. U.D. conceptualized the work and interpreted the data. S.P.P. conceptualized the work and interpreted the data. S.J.P. conceptualized the work and interpreted the data. J.M.D. conceptualized the work and interpreted the data. All authors revised the paper.

## Competing Interest Statement

Joseph M DeSimone reports a relationship with Carbon that includes board membership and cofounder equity. Provisional patents have been filed by Stanford University for the reported devices.

## Materials and Methods

### POP Device Designs and Fabrication

Three versions of the POP device were used: version A, version B, and version C (Fig. S1). All three versions have the same basic design, with A and B differing only in the MAP used, and C being a larger design with additional accessories for use in the clinical study. All components were designed in Fusion 360 (Autodesk, San Francisco, CA).

Version A was used to study spatial pressure distribution in *ex vivo* human skin. It consisted of a microneedle array patch (MAP) and a perforated collection plate affixed to a vacuum adapter. The MAP featured a 7 × 5 array of tapered hollow microneedles, with a base diameter of 550 µm tapering to 400 µm at the tip, and a 200 µm wide hollow channel. Each microneedle was 2200 µm long, and the tip formed a 55° angle to the tapered wall. Pitch was 1000 µm. To prevent overcure, the center three microneedles were enlarged to 450 µm at the tip to allow a 250 µm wide channel. The MAP was made from KeySplint Hard resin (Keyprint #4220004) (Fig. S2).

The collection plate measured 9 × 6 mm and had a 7 × 5 array of perforations, each 500 µm in diameter and 700 µm deep (Fig. S2); seven central perforations were enlarged to 600 µm in diameter to mitigate overcure artifacts during 3D printing. The MAP-facing side of the plate featured 400 µm tall spacer pegs such that 1100 µm of the microneedles were exposed when mated with the plate. The plate was made from KeySplint Soft Clear Resin (Keyprint #4220161). The plate was assembled with a custom vacuum adapter so that a facial vacuum equipped with a 20x magnification camera (Prabensei Model TMH100, Amazon) can be attached to apply vacuum and visually monitor ISF extraction in real-time. Together, the assembled unit is referred to as the collection apparatus. The vacuum adapter was made from either KeySplint Soft Clear (Keyprint #4220161) or KeySplint Soft Resin (Keyprint #4220005), with no functional difference. For the pressure ring variant of version A (Fig. S3), ring protrusions with 140 µm thick walls and 500 µm height were added to the skin-facing side of the plate. The MAP featured 500 µm base diameter hollow microneedles that were 2700 µm in length to fit the modified plate.

Version B was used to study penetration depth, collection time, and pressure (Fig. S2). This version was identical to version A except the MAP was replaced with a solid needle design with 1600 µm tall cylindrical posts with 600 µm tall conical tips. Diameter of the microneedle was 400 µm. The MAP was made of KeySplint Soft Clear resin (Keyprint #4220161), which has a reduced fracture rate compared to KeySplint Hard resin.

Version C (clinical) had the same basic design as version B, with modifications to expand the MAP and collection plate array size to 154, a bracket made of KeySplint Hard for assembling the collection plates and for handling, a topper made of KeySplint Soft that replaced the MAP after MAP removal to ensure uniform pressure application, and a stand made of KeySplint Hard for positioning the spring-loaded applicator at a controlled height (Fig. S10). No vacuum adapter was used in Version C.

All POP device components were 3D printed using either a prototype S2 printer with 5.4 µm lateral resolution or an M2 printer with 75 µm lateral resolution from Carbon 3D. The MAPs and collection plates were printed using the S2 while all other components and accessories were printed using the M2. For MAPs, dynamic printing in step mode was used, with slice thicknesses of 25-100 µm, exposure times of 0.9-3 s, and 25 mW/cm^2^ UV intensity at 385 nm wavelength. Print time varied from 7 to 11.5 minutes. 1-3 MAPs fitted on the build platform. For plates, the print parameters were 25 µm slice thickness, 0.3 s exposure time, and 25 mW/cm^2^ UV intensity at 385 nm wavelength. For parts printed on the M2, built-in print parameters for each resin were used. POP production for the clinical study took place in a workspace equipped with build platforms and vats dedicated to human-grade parts.

After printing, a part was gently removed from the build platform with a razor and sonicated in 100% isopropanol (IPA) for 30 seconds twice, cleaning it with pressurized air after each sonication. The part was then cured for 32 minutes at 90% intensity in nitrogen in a Dentamid PCU LED N2 UV oven (Dreve #4317). Collection plates were cured for 32 minutes at 90% intensity in air since oxygen removal during curing was found to reduce the plate’s hydrophilicity.

The collection plate was assembled to the vacuum adapter or bracket by using Keysplint Hard as a glue and curing for 32 minutes at 90% intensity in air. The assembled part was sonicated in IPA for 30 seconds twice, cleaning with pressurized air after each sonication.

For human use, a set of two MAPs and a stand were sealed in a pouch and sterilized in an autoclave. A pre-vacuum sterilization cycle was used, with 250°F temperature, 30 minutes sterilization time, and 5 minutes drying time.

### Porcine and Human Skin Sources

Porcine ear skin was harvested from recently euthanized Yorkshire pigs following methods adapted from Golombek et al^34^. The outer ear skin was shaved and dissected from cartilage using a scalpel. The isolated skin was briefly sterilized in a mixture of Povidone-Iodine solution (Betadine) and PBS. The skin was then washed in PBS, patted dry with a Kimwipe, and stored at -20°C.

Frozen food grade porcine back skin was purchased from an online retailer that sourced from Pacific Agri Products, Inc. (South San Francisco, CA). The skin was stored at -20°C.

Flash-frozen human skins from the abdomen were purchased from ZenBio (ZenBio T-SKN-FF10CM) and the National Disease Research Interchange (NDRI). The NDRI flash freezing protocol involved transport on dry ice immediately after harvest and storage at -80°C. The samples were stored according to vendor guidelines. Skins from ZenBio were stored at -20°C. Skins from NDRI were stored at -80°C. Portioned pieces were transferred from -80°C to -20°C in batches for use to prevent degradation from freeze-thaw cycles.

### ISF Collection in Skin Ex Vivo

An approximately 2 cm × 1.5 cm piece of human (Table S4) or porcine skin was cut from the frozen sample and thawed at room temperature in a petri dish. Subcutaneous fat, if present, was trimmed using a razor. After thawing, the skin was thoroughly patted dry using a Kimwipe and transferred onto a piece of parafilm on a hard benchtop. The hardness of the surface can influence MAP penetration into the skin.

The collection apparatus was weighed with an analytical balance (Mettler Toledo, XSR105) to record the pre-collection dry weight (Fig. S24). A MAP was mated with the collection apparatus, and the complete assembly was placed on the skin. A commercial spring-loaded applicator (MPatch Micro Applicator, Micropoint Technologies) with a custom extender (Fig. S25) was positioned on the collection apparatus. The applicator was deployed three times to strike the MAP.

After skin insertion, the MAP was carefully removed with tweezers without shifting the collection apparatus. Misaligned punctures with collection plate perforations negatively impact ISF collection. The MAP was inspected for fractures post-insertion. The entire setup was transferred with parafilm onto a silicone pad (suture practice pad, ASIN B0842L4T44, Amazon), used to simulate subcutaneous fat. In our experience, ISF collection is equally effective on a hard surface. A 500 g weight (calibration weight set, HFS, CKW-1000G, Amazon) was placed on the collection apparatus to apply pressure for 5 minutes, balanced by hand to prevent toppling. After removing the weight, the collection apparatus was slowly peeled away from the skin. Operator variability was observed during this step (Fig. S26). If desired, a 20x camera can be installed on the collection apparatus to visually monitor the removal process. Surface tension held the collected ISF in the plate. A post-collection weight was recorded to determine the ISF amount.

After collection, the skin was stained topically with 20-40 µL of 2% w/v methylene blue dye in PBS for 5 minutes before rinsing thoroughly with water. This step dyed the puncture sites where the stratum corneum was breached. The skin was then imaged under a microscope (Olympus DSX1000, Evident Scientific, San Jose, CA) to ensure each microneedle on the patch punctured successfully (Fig. S27). Occasionally, one or two punctures may fail. Skin staining was omitted once a device’s puncture efficiency was confirmed more than a few dozen times.

For experiments requiring isolation of ISF from POP, the collection apparatus containing ISF was fitted on a 50 mL conical tube using parafilm. 100 µL of diluent, typically PBS, was pipetted onto the collection plate to increase handling volume. The diluted ISF was collected in the tube after centrifugation at 2000 G × 10 s (ThermoFisher Scientific Sorvall XR Pro).

### Pressure Distribution Experiments

Experiments used version A of the POP device. Skin sourced from a single donor was used to minimize biological variability. Methods described in the ISF Collection section were followed using *ex vivo* human skin. A modification was made to apply pressure on a bench top rather than on a silicone pad. The weight used was 500 g. Collection time was 5 min. For the POP scenario, standard procedures were followed. For the needle-in scenario, MAP was not removed. An insert was placed on top of the MAP (Fig. S25), and the weight was placed on the insert to apply pressure to the MAP and collection apparatus for the duration of 5 minutes. The MAP and collection apparatus were removed together at the end of the collection. For the vacuum scenario, instead of placing a weight on top of the collection apparatus, a vacuum (Prabensei Model TMH100, Amazon) was attached and set to high (−64 kPa, Fig. S28) for 5 minutes. For the pressure ring scenario, the standard procedure was followed using a collection plate modified with ring protrusions. For the control, no pressure was applied during the 5 minute period. A custom insert (Fig. S25) was placed in the collection apparatus to minimize evaporation.

### Finite Element Analysis

Fusion 360 (Autodesk, San Francisco, CA) was used for finite element analysis (FEA). Only the dermis was simulated using a simplified one-layer elastic model (Table S5), measuring 3.5 × 3.5 mm laterally and 1.5 mm in thickness. A collection plate measuring 3 × 3 × 0.5 mm with a 3 × 3 array of 500 µm diameter perforations was placed on the dermis. The plate was simulated as steel to approximate a rigid body. For the POP scenario, a force equivalent of 53.9 kPa (0.39 N) was applied to the plate. For the needle-in scenario, the plate was attached to a solid MAP with 500 µm tall and 200 µm diameter conical tips. Puncture sites were modeled in the dermis to match the MAP. 0.39 N of force was applied to the MAP and plate as a single body. This approach simplified the simulation over using a complex hollow needle geometry, without altering the qualitative features of the resulting pressure distribution. For the vacuum scenario, a force equivalent of -53.9 kPa (−0.094 N) was applied to each skin site matching a plate perforation. For the pressure ring scenario, the plate was modified with 500 µm tall ring protrusions with 140 µm thick walls. 0.39 N of force was applied to the plate. For display, the ZZ component of the stress tensor was shown. A scale from -40 kPa (compression) to 10 kPa (tension) was used except in the vacuum scenario, where the scale was 15 kPa to 65 kPa because the dermis was mainly in tension. Since ∇p rather than absolute p determines ISF flow, the different scenarios can be compared as long as the display scale range stayed consistent. Only the center of the array is shown, as each perforation can be treated as a repeating unit.

### Penetration Depth Experiments

Experiments used version B of the POP device. Penetration depth was measured on porcine skin because samples from different anatomical regions were available. First, a MAP was mated with a collection plate and placed on the skin, without a vacuum adapter. Then, a vacuum adapter was positioned over the MAP and the plate to recreate the penetration conditions used for ISF sampling. A spring-loaded applicator was positioned on the adapter and deployed three times to strike the MAP. After removing the applicator and vacuum adapter, the MAP and plate were secured to the skin with Tegaderm. The skin was fixed overnight in 5 mL of 10% formalin. The next day, the MAP and plate were removed, and an optical coherence tomography (OCT) (Ganymede SD-OCT, ThorLabs) scan was performed. OCT slices were analyzed with ImageJ to quantify penetration depth at each puncture site. Every puncture site could be measured by this method.

For cross-validation, penetration depth was also measured by optical microscopy. A MAP was dipped in tattoo ink (Dynamic Color Co., black, Amazon) to coat the tips, and applied to the skin using the method described above. After overnight fixation and MAP removal, the skin was sliced with a razor along a row of puncture sites using tattoo ink as visual guidance. The tissue slices were placed between glass slides and imaged with an Olympus DSX1000 (Evident Scientific, San Jose, CA). Penetration depth was quantified from the ink staining using ImageJ. Only a fraction of the puncture sites could be measured by this method due to slicing limitations.

Histological slides were prepared by Histo-Tec Laboratory (Hayward, CA). Samples were submitted in 10% formalin after overnight fixation. Paraffin embedding, sectioning, and hematoxylin and eosin (H&E) staining were performed. Tattoo ink aided in distinguishing puncture sites from nonspecific cuts and tears.

To measure ISF collection, methods described in the ISF Collection section were followed. A modification was made to first immerse the back skin in PBS for 5 minutes to ensure a similar hydration level to the ear skin. For the control experiment, ISF was collected from the underside of the skin without MAP puncture, directly from the ISF-rich region of the dermis.

### Collection Time and Pressure Experiments

Experiments used version B of the POP device. To minimize operator variability during collection apparatus removal from the skin (Fig. S26), a gauze pad (HealQU, Non-woven 2”x2”, ASIN B0BL46LXRN, Amazon) was cut to size and adhered by friction to the skin-facing side of the collection apparatus. This gauze retained the collected ISF instead of the plate perforations, preventing partial ISF loss during plate removal. After the gauze is adhered, a MAP is mated with the plate, and tweezers were used to push down the gauze around each microneedle to expose them (Fig. S24).

To measure ISF collection, methods from the ISF Collection section were followed using *ex vivo* human skin with varying times and pressures. Skin from a single donor was used for each time and pressure series to reduce biological variability. Time points were 0, 1, 5, 15, and 30 minutes at a constant pressure of 50 kPa. Calibrated weights from 200 to 1000 g were used to vary pressure at a constant time of 5 minutes. For weights over 500 g, a custom 3D-printed holder was used to support multiple weights (Fig. S25). The weight of the holder was included in the pressure calculations. For the zero pressure experiment, a custom insert (Fig. S25) was placed in the collection apparatus to minimize evaporation. The weight of the insert equaled 0.9 kPa of pressure. At the end of collection, the gauze holding ISF may stick to the skin. In these cases, the gauze was manually transferred to the collection apparatus with tweezers before weight measurement. After weighing, the gauze was immersed in 100 µL of 200 mM ammonium bicarbonate buffer in a microcentrifuge tube to extract the ISF. ISF protein concentration was quantified, and the sample was stored at -80°C. For volcano plot analysis, comparisons include 1 min (N=6) and 30 min (N=9) samples collected at 50 kPa, 0.9 kPa (N=9) and 104 kPa (N=9) samples collected at 5 min, and control samples collected without pressure at 0 min (N=12) and 30 min (N=3). The two sets of control samples differ only by degree of evaporation. Replicates represent three mass spectrometry injections for each biological replicate. Due to the minuscule ISF collected without pressure or at short time points, some samples lacked sufficient proteins for reliable MS analysis, resulting in missing replicates.

### Human Study

The study used version C (clinical) of the POP device. All procedures followed aseptic techniques. 10 subjects participated in the study. 5 subjects additionally underwent finger prick blood collection on the same arm as ISF collection. All subjects provided informed consent. Separately, 3 subjects underwent sham collections without MAP puncture.

For blood collection, a finger was cleaned with an alcohol pad, then pricked with a OneTouch Delica Plus 33G lancet at the deepest setting (setting = 7). Blood was collected using a calibrated capillary tube marked in 1 µL increments (Cat #2-000-001, Drummond Scientific, Broomall, PA). A photo was taken to measure the volume collected to a precision of 0.1 µL via analysis in ImageJ. The blood was expelled into 100 µL of PBS using an aspirator and placed on ice.

For ISF collection, the collection plate and topper were weighed beforehand. The procedure was performed on the volar forearm. A warm pack (HotHands Hand Warmers, Amazon) was applied for 5 minutes to increase ISF volume. The efficacy of warming was not studied. During this time, the MAP was removed from packaging and mated to the collection plate. After cleaning the skin with an alcohol pad, the assembled MAP and plate were placed on the arm. The applicator stand was positioned around the assembly (Fig. S10). A commercial spring-loaded applicator modified with a 25 N spring (PC031-281-7500-MW-0599-CG-N-IN, The Spring Store) was placed on the stand and deployed three times, each time sliding into a different predefined location on the stand to evenly strike the entire MAP surface.

After skin puncture, the MAP was removed carefully, keeping the collection plate still. A topper was placed on the plate. A thin-film force sensor (Qinlorgo, RP-C-MK01X, Amazon) was placed on the topper, and 750 g of thumb pressure, equivalent to 25 kPa, was applied for 5 minutes. The plate and topper were carefully removed in a peeling motion and weighed to measure the ISF collected. The skin site was photographed, cleaned with an alcohol pad, and dressed with an adhesive bandage. Pain was rated on the Visual Analog Scale.

After ISF collection, the collection plate and topper were gently detached. 100 µL of PBS was pipetted onto the topper, and the plate was re-attached. PBS should distribute evenly on the plate to dilute the ISF. The whole unit was placed face down in a custom adapter (Fig. S25) fitted on a 15 mL conical tube and centrifuged at 2000 G × 10 s. Post-centrifugation, ISF was extracted into the tube with 80% volume recovery, while RBCs settled at the bottom. The top and bottom halves were pipetted into separate tubes. The top half, labeled ISF supernatant, was contaminated only by hemolyzed RBCs. The bottom half, labeled ISF sediment, was contaminated by both hemolyzed and intact RBCs. Protein concentration and blood contamination were measured, and the remainder was stored at -80°C. Measurements were not possible for one subject due to sample mishandling resulting in unknown ISF dilution ratio.

After blood collection, hemoglobin and RBC counts were measured. Then, the sample was centrifuged at 200 G × 5 min at 4°C to form an RBC pellet. The supernatant was transferred to a microcentrifuge tube and labeled as plasma. The pellet was resuspended in 100 µL of 200 mM ammonium bicarbonate and labeled as RBCs. RBCs were lysed in the ammonium bicarbonate buffer. Protein concentration was measured in RBC. Protein concentration and blood contamination were measured in plasma. The remainder was stored at -80°C.

### Protein Concentration Measurement

Protein concentrations were measured using UV-vis absorbance and BCA assay for cross-validation. The mean across the two methods was reported. A few samples lacked sufficient volume for both measurements, so only UV-vis absorbance was measured. For samples with intact RBCs, such as ISF sediment, the sample was diluted 10-fold in 200 mM ammonium bicarbonate, vortexed, and left for 10 minutes to ensure hemolysis before UV-vis measurement. For pure RBCs, only UV-vis absorbance was used because the BCA assay underestimated protein concentration when crossvalidated against hemoglobin concentration, possibly due to redox interference by heme.

UV-vis absorbance was measured with a NanoDrop One (Thermo Fisher Scientific) using the Protein A205 method based on peptide bond absorbance. Measurements assumed an extinction coefficient of 31 cm−^1^mg−^1^ml at 205 nm without baseline correction to avoid errors from hemoglobin. For experiments utilizing gauze, the contribution of bare gauze to the 205 nm absorbance, about 3-4% on average, was subtracted. BCA assays (Thermo Fisher Scientific) were conducted per the manufacturer’s instructions using bovine serum albumin as the standard, with absorbance measured at 560 nm. Standard curve was fitted to a quadratic model. Hemoglobin was subtracted from total protein in ISF and plasma to reduce errors from blood contamination. Plasma volume was assumed to be 55% of whole blood volume when calculating plasma protein concentration.

### Blood Contamination Measurement

Blood contamination was estimated by both UV-vis absorbance and RBC count for cross-validation. Both data points were presented for each subject. For ISF supernatant, only UV-vis absorbance was used due to the absence of RBCs. For ISF sediment containing RBCs, the sample was diluted 10-fold in 200 mM ammonium bicarbonate, vortexed, and left for 10 minutes to ensure hemolysis before UV-vis measurement.

UV-vis absorbance was measured with a NanoDrop One (Thermo Fisher Scientific) using a custom oxyhemoglobin method by the manufacturer. Hemoglobin was assumed to be 100% oxygenated, compared to the typical 95-98% in blood. The oxyhemoglobin concentration was manually calculated using a molar extinction coefficient of 524,280 M^− 1^cm^− 1^, a molecular weight of 64,500 g/mol, and the raw absorbance at 414 nm^35^. For RBC count, a hemocytometer was used after approximately 1:200 dilution of whole blood in PBS. Blood contamination was estimated by calculating ISF hemoglobin and RBC count as fractions of those in whole blood. If blood was not sampled, normal values of 150 mg/mL for hemoglobin and 5 million cells/µL for RBC count were assumed. Contamination in ISF immediately post-collection was calculated from measured contaminations in ISF supernatant and ISF sediment, and their known respective fractions. Calculation of ISF contamination by RBC count accounted for hemolysis, as hemolyzed cells would be excluded from the count.

Data from 6 of the 10 subjects was presented for blood contamination. In the first 3 subjects, an incomplete post-processing protocol caused total hemolysis and a lack of separation between ISF supernatant and sediment. One sample was mishandled, resulting in an unknown ISF dilution factor after PBS addition. Among the 6 subjects, 4 underwent simultaneous blood collection.

### Mass Spectrometry

Samples were kept on ice during handling. All buffers and solvents used were LC-MS grade. For protein digestion, 25 µg of protein was aliquoted, and brought up to 50 µL of final volume with 50 mM ammonium bicarbonate. The ammonium bicarbonate buffer was made fresh daily. Then, 2.5 µL of 200 mM Tris(2-carboxyethyl) phosphine (TCEP) (Sigma-Aldrich) was added for disulfide reduction, followed by a 1-hour incubation at 65°C. After incubation and allowing the samples to reach room temperature, 3.75 µL of freshly prepared 200 mM Iodoacetamide (Sigma-Aldrich) was added. The sample was covered in aluminum foil and incubated for 45 minutes at room temperature. 1 µg of sequencing grade modified trypsin enzyme (Thermo Fisher Scientific) prepared at 0.2 µg/µL in 50 mM ammonium bicarbonate buffer was added. After vortex mixing, the sample was incubated overnight at 37°C. After digestion, the sample was dried using a speed vacuum (LabConco).

For samples diluted in PBS, desalting was performed following digestion using C18 ZipTips with 0.6 µL beads and 10 µL volume (Millipore). Wetting buffer was 0.1% formic acid in 100% Acetonitrile. Equilibration buffer was 0.1% formic acid in water. Wash buffer was the same as the equilibration buffer. Elution Buffer was 0.1% formic acid in 80% acetonitrile and 20% water. To begin, dried samples were reconstituted in 13 µL of equilibration buffer. Next, microcentrifuge tubes were arranged in a specified order: Wetting, Equilibration, Sample, Wash 1, Wash 2, and Elute. 40 µL of the appropriate buffer is added to each tube. With the pipettor volume set to 10 µL, solution was aspirated and dispensed into the tube through the ZipTip 10 times at each step. The process was then repeated starting at the equilibration step to increase yield. The eluted peptides were dried using a speed vacuum (LabConco).

For mass spectrometry analysis, the digested sample was reconstituted in 0.1% formic acid in water, and approximately 1 µg of tryptic peptides was injected into a 10 µL injection loop in tandem to Dionex Ultimate Rapid Separation liquid chromatography system (Thermo Fisher Scientific). The sample was loaded onto a PepMap Neo C18 trap column (Thermo Fisher Scientific) at a constant flow rate of 5 µL/min for 10 minutes with 2% (v/v) acetonitrile and 0.1% formic acid in LC/MS grade water. Eluted peptides were resolved on a 25 cm long analytical column packed with BEH C18 1.7 µm particles (Waters) by reversed-phase liquid chromatography. A gradient consisting of mobile phase A (0.1% (v/v) formic acid in LC/MS grade water) and mobile phase B (0.1% (v/v) formic acid in acetonitrile) was used to elute tryptic peptides from the analytical column, by holding mobile phase B at 2% for the first 6 minutes, ramping up from 2% to 35% mobile phase B over the next 125 minutes, and ending with an increase to 85% mobile phase B over 5 minutes with a 5-minute hold at a constant flow rate of 0.3 µL/min. A 10-minute equilibration time with 2% mobile phase B was added at end of the gradient prior to the next sample injection. Each sample was injected in triplicates. A column heater (PST Phoenix S&T) was used to heat the column to a temperature of 65 °C. Eluted peptides underwent MS/MS analysis on an Orbitrap Tribrid Eclipse mass spectrometer (Thermo Fisher Scientific). 2.2 kV was applied to a nano-spray source (Thermo Fisher Scientific). The ion transfer tube temperature was 275 °C. The Orbitrap MS1 mass resolution was set to 240,000 with a scan range of 375 - 1800 m/z. The radio frequency (RF) was set to 30%, with a max injection time of 35 ms and ion intensity threshold of 5.0 × 10^3^ per MS1 scan. MS2 scans were acquired using Top-Speed with a cycle time of 1 ms. Dynamic exclusion was enabled for 30 s with a mass tolerance of 10 ppm. Ion precursor isolation was performed in the quadrupole with a mass isolation window of 0.7 m/z. Ion precursors were subject to higher-energy collisional disassociation (HCD) with a fixed collisional energy of 28%. Fragmented ions were detected in the linear ion-trap using turbo scan rate. The automatic gain control (AGC) target value was set to standard with a maximum injection time of 50 ms.

### Proteomic Data Analysis

MaxQuant software (version 2.6.3, Cox Group, Max Planck Institute) was used for protein identification, intensity-based absolute quantification (iBAQ)^27^, and label-free quantification by the MaxLFQ algorithm^36^. The FASTA file for protein identification was sourced from UniProt, containing approximately 20,000 Swiss-Prot human protein sequences, accessed on July 24, 2024. Default parameters were used, except for the following: LFQ minimum ratio count was set to 1, fast LFQ was disabled, match between runs was enabled, and iBAQ was enabled. Common contaminant proteins identified in the FASTA file were removed from analysis, except for albumin, gelsolin, periostin, dermokine, metallothioneins, tropomyosin, osteopontin, tetranectin, and thymosin beta-10. The retained proteins included well-established plasma proteins, skin proteins that may be found in dermal ISF, or those with no clear reason for contaminant classification. Proteins identified by a single modification site were excluded from the analysis.

Statistical analyses were performed using R (version 2024.09.0, CRAN). T-tests performed were two-sided assuming heteroscedasticity. For plasma and ISF comparisons from the same subject, paired t-tests were performed, and the resulting p-value distribution was used to calculate Storey’s q-value^37^. Parameter λ in Storey’s procedure was selected automatically or manually to match a stable π_0_ region.

To generate a ranked list of RBC contaminant proteins from most to least contaminating, we ranked proteins by mean Log_2_(LFQ RBC) - Log_2_(LFQ ISF) across 5 pairs from 5 subjects (Fig. S16, S29). Only proteins with at least 2 pairs and an FDR < 0.05 were included. Next, proteins exclusively detected in RBC, defined as detectable in >66% of RBC MS runs and <33% of ISF MS runs, were inserted into the ranked list. These proteins represent additional potential contaminants not readily detected in ISF samples at <1% blood contamination. The insertion position was based on the mean LFQ RBC for each protein since the RBC-ISF difference cannot be calculated. A similar ranked list of plasma contaminant proteins was generated (Fig. S17, S30). IGLC2, IGLC3, and IGLC6 were removed from the plasma contaminant proteins list due to their high degree of homology, which can lead to artificial discrepancies in ISF and plasma levels caused by peptide assignment errors.

### Statistical RBC Contamination Removal

The volume percentage of blood contamination in ISF was estimated by hemoglobin following methods described in the Blood Contamination Measurement section. Next, the percentage and number (N) of contaminated proteins in the proteome, defined as >10% quantification error resulting from contamination (Fig. S15), was calculated. Finally, the ranked list of RBC contaminant proteins was traversed, removing each contaminant from the ISF proteome until N is reached. If a contaminant protein was absent in the ISF sample, the protein was skipped and the list traversal continued. A less stringent definition of contaminated protein, such as >20% quantification error, can be selected depending on the research question.

### Ethics Statement

All experiments involving *ex vivo* animal tissues were approved by the Institutional Animal Care and Use Committee (IACUC) and the Administrative Panel on Laboratory Animal Care (APLAC) at Stanford University. Porcine ear skins were harvested from euthanized Yorkshire pigs per APLAC protocol #31492.

For human skins sourced from ZenBio, consent was obtained under IRB oversight by the vendor. For human skins sourced from the NDRI, an IRB exemption was obtained from Stanford University. The pilot human study was conducted at Stanford University under IRB protocol #74098. Inclusion criteria include 18 years of age or older. Exclusion criteria include history of skin conditions that predispose the participant to adverse reactions to skin puncture, active infection, open wound, rash, or skin breakdown on the arm being used for MAP application, and known acrylate allergy.

### Software for Figure Preparation

Figures were produced using R (version 2024.09.0), Inkscape (version 1.4), Gimp (version 2.10), BioRender, PowerPoint (Microsoft 365), and GraphPad Prism (version 10.2.2).

