## Supplementary Information for "A Microneedle Device for Rapid Dermal Interstitial Fluid Sampling"

Andy H. Hung[1,7], Netra U. Rajesh[2,7], Abel Bermudez[1,3], Stephanie M. Boczek[5], Fernando J. Garcia-Marqués[1,3], Yee Lin Tan[6], Jihyun (Luna) Hwang[5], Prima Dewi Sinawang[3,5], Dan Ilyin[4], Gunilla B. Jacobson[1], Utkan Demirci[1,3], Steven P. Poplack[1], Sharon J. Pitteri[1,3], and Joseph M. DeSimone[1,3,5,*]

[1] Department of Radiology, School of Medicine, Stanford University, CA 94305

[2] Department of Bioengineering, Stanford University, CA 94305

[3] Canary Center, School of Medicine, Stanford University, CA 94305

[4] Department of Mechanical Engineering, Stanford University, CA 94305

[5] Department of Chemical Engineering, Stanford University, CA 94305

[6] Department of Materials Science and Engineering, Stanford University, CA 94305

[7] These authors contributed equally to this work

**Table of Contents**

**1. Supplementary Figures**

S1. Overview of different POP variants
S2. Dimensions and micrographs of versions A and B of POP used in *ex vivo* experiments
S3. Modified version A of POP for applying pressure in ring distributions
S4. Comparison of solid and hollow MAPs in POP
S5. Comparison of vacuum application after MAP puncture in human skin *ex vivo*
S6. Skins used in *ex vivo* experiments
S7. Cross-validation of penetration depth measurements
S8. Control experiment for penetration depth study
S9. Protein molecular weight and ISF collection pressure
S10. Applicator stand for POP
S11. Mechanical testing of solid MAPs
S12. Characterization of monomer leaching in POP
S13. Photographs of POP puncture sites in human subjects
S14. ISF collection after skin pre-treatment with hyaluronidase
S15. Distribution of mass spectrometry quantification coefficient of variation
S16. Ranked list of RBC contaminant proteins
S17. Ranked list of plasma contaminant proteins
S18. Nanoparticle tracking analysis (NTA) of human ISF
S19. Comparison of the most abundant proteins in human plasma and ISF
S20. Potential protein biomarkers in ISF
S21. Underestimation of large molecule concentration in ISF due to volume exclusion
S22. Microcentrifuge adapter for ISF extraction from collection plate
S23. Monolithic latticed MAP device
S24. ISF collection procedure in skin *ex vivo*
S25. ISF collection by POP varied by operator technique
S26. Examples of MAP skin puncture verification by methylene blue dye
S27. Custom 3D-printed experimental accessories
S28. Pressure measurement for commercial vacuum
S29. RBC contaminant proteins visualized within the RBC proteome by abundance
S30. Plasma contaminant proteins visualized within the plasma proteome by abundance

**2. Supplementary Tables**

S1. Demographics in human study
S2. Negative control to rule out ISF contamination by skin surface proteins
S3. Comparison of immunoglobulin quantification with previous work
S4. Demographics of human skin used for *ex vivo* experiments
S5. Finite element analysis skin model parameters

**3. Supplementary Methods**

A. Hyaluronidase Skin Pretreatment
B. Comparison of Solid and Hollow MAPs in POP ISF Collection
C. Mechanical Testing

**4. Supplementary References**

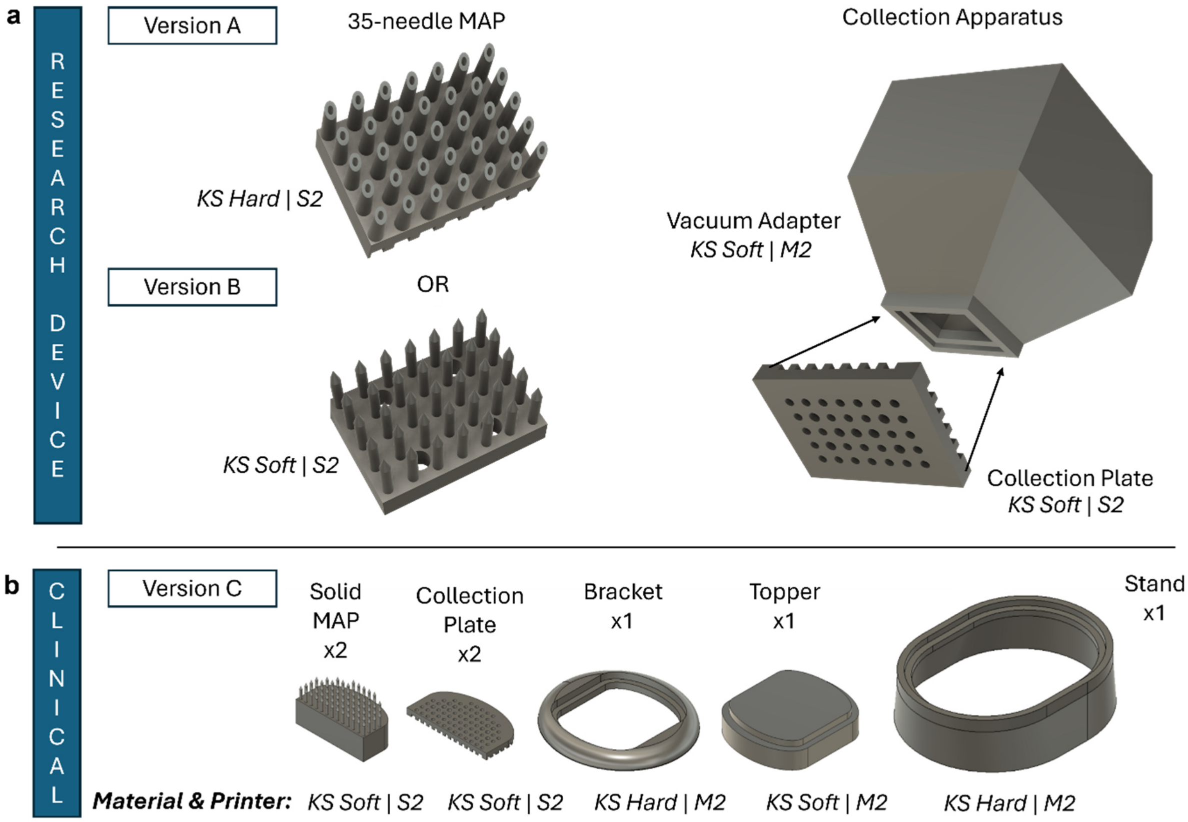


**Figure S1. Overview of different POP variants.** Version A was used in the pressure distribution study. Version B was used in the time series, pressure series, and penetration depth studies. Version C was used in the human study. **a** Versions A and B differ only in the MAP component, with a hollow design in A and a solid design in B. The collection plate is attached to a vacuum adapter by resin cure for vacuum application and video monitoring. The assembled unit is referred to as the collection apparatus. **b** Components of the clinical POP device. The resin material and Carbon 3D printer used are denoted as “resin material | printer”.

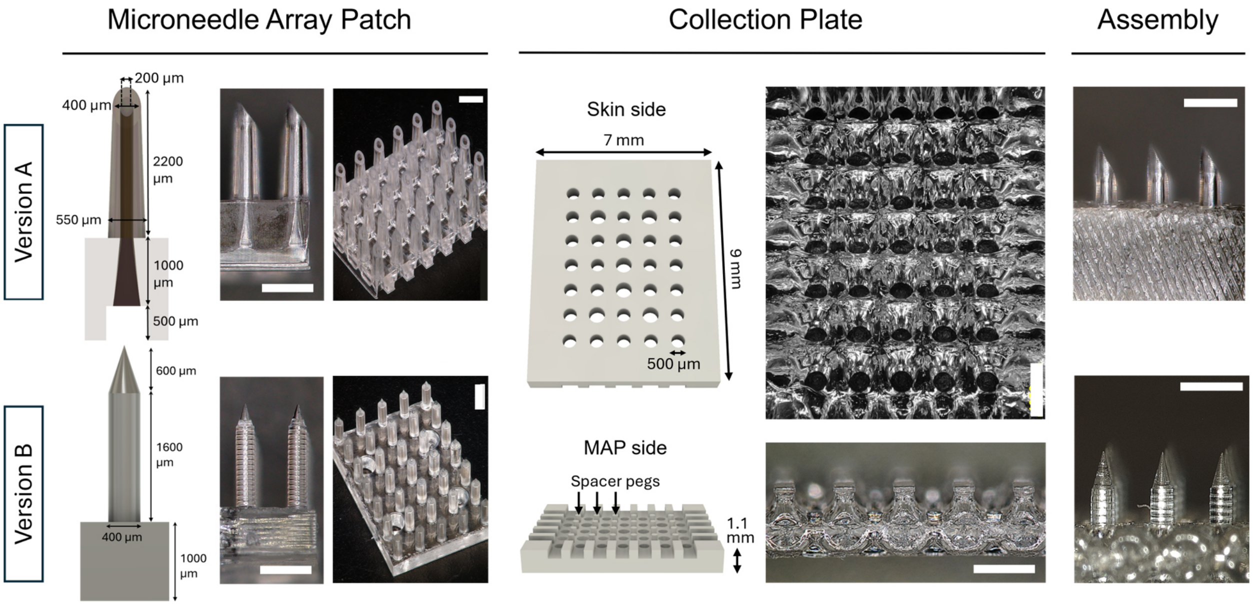


**Figure S2. Dimensions and micrographs of versions A and B of the POP device used in ex vivo experiments.** The two are identical except for the MAP component. Surface ripples on the collection plate are from 3D printing artifact with Keysplint Soft. Central perforations on the collection plate are slightly enlarged to 600 µm to compensate for overcure. Scale bars = 1 mm.

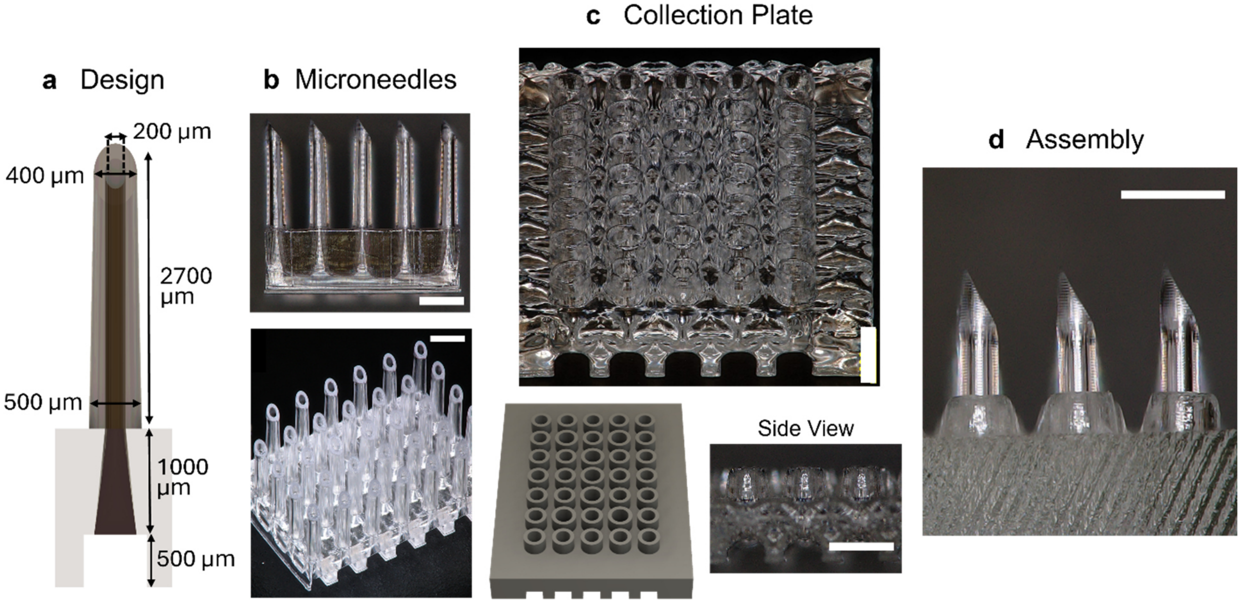


**Figure S3. Modified version A of POP for applying pressure in ring distributions. a** Dimensions of the microneedle, lengthened to accommodate the height of the rings on the collection plate. **b** Micrographs of microneedles. **c** Collection plate modified with ring protrusions featuring 140 µm thick walls and 500 µm height. **d** Micrograph showing the exposed microneedle length after mating with the collection plate. Scale bars = 1 mm.

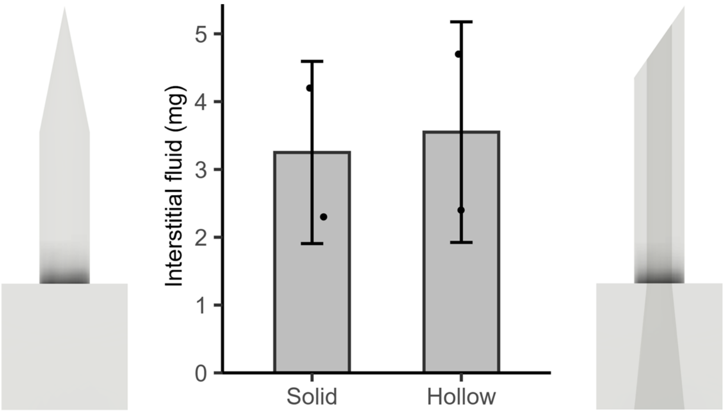


**Figure S4.** POP collects similar amounts of ISF from human skin *ex vivo* using either a solid or a hollow MAP for puncture.

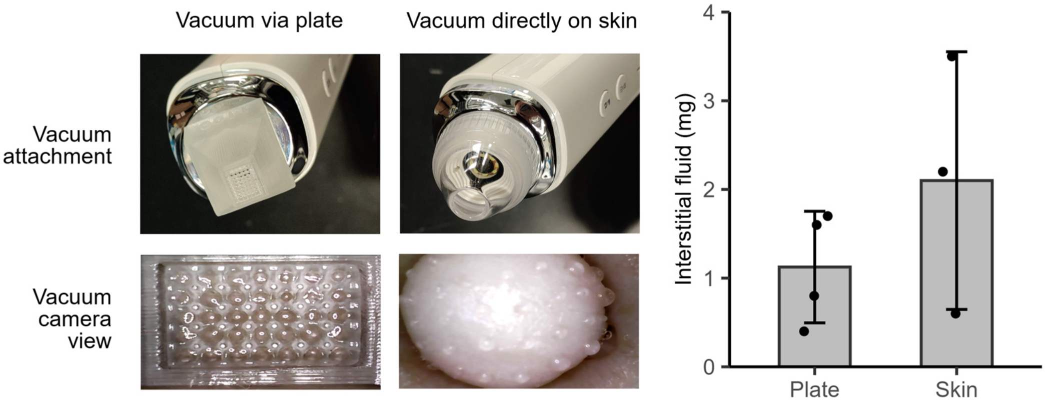


**Figure S5. Comparison of vacuum application after MAP puncture in human skin *ex vivo*.** By switching attachments, vacuum can be applied to the skin via a collection plate or directly after MAP puncture. Sustained vacuum seals around all 35 openings were likely not achieved with the collection plate, resulting in suboptimal collection. In contrast, applying vacuum directly to the skin secured an excellent seal, confirmed by skin dome formation and ISF droplets at the puncture sites. However, the pressure gradient in the lateral dimensions was eliminated. ISF collection sometimes approached that of POP in this configuration. -64 kPa was applied for 5 minutes.

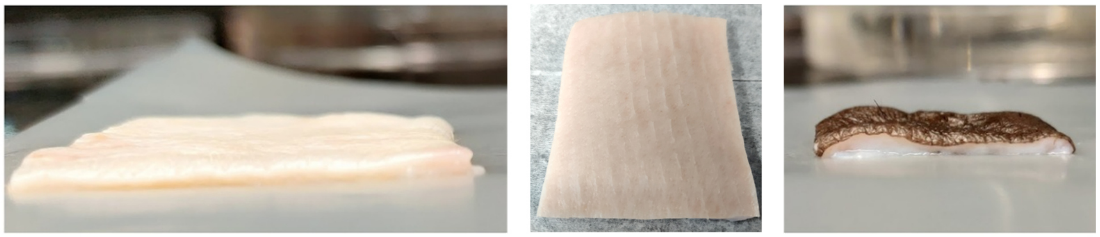

**Figure S6. Skins used in *ex vivo* experiments.** Left: porcine ear skin (1.6 mm thickness). Middle: porcine back skin (3.5 mm thickness). Right: human abdomen skin (1.5 mm thickness).

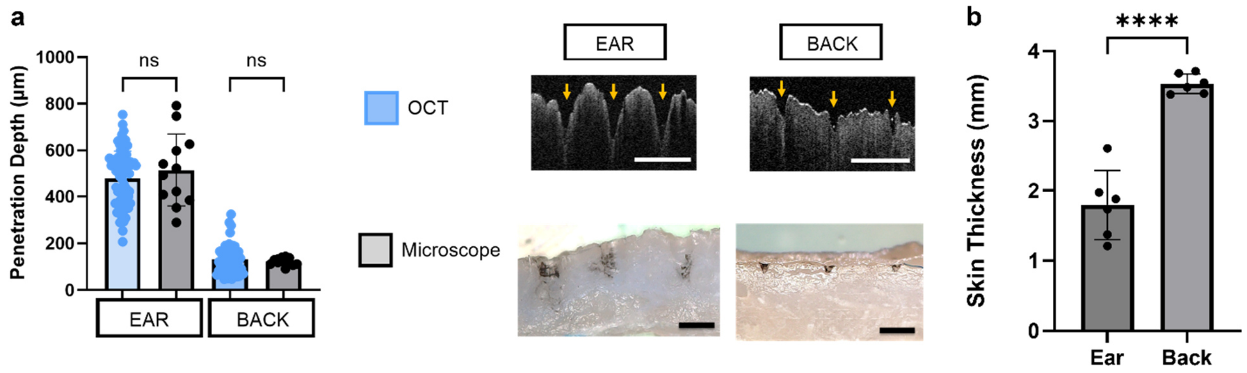


**Figure S7. Cross-validation of penetration depth measurements. a** Comparison of penetration depth in porcine ear and back skin measured using optical coherence tomography (OCT) and optical microscopy. Representative images show the penetration sites. Good agreement was achieved between the two measurement methods. Scale bars = 500 μm. **b** Thickness of porcine ear and back skin samples measured by digital calipers (N = 6). Statistical analysis used a two-way ANOVA with post-hoc Tukey test. **** = $p < 0.0001$. ns = not significant.

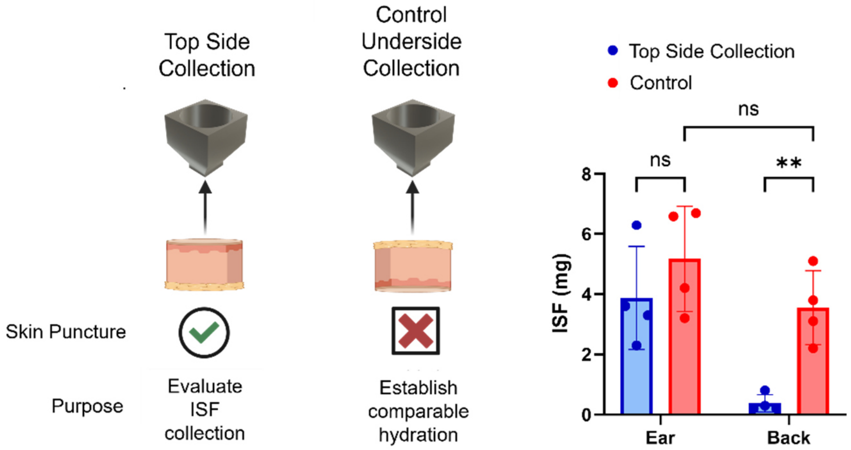


**Figure S8. Control experiment for penetration depth study.** To verify that ISF collection differences between porcine ear and back skin were due to penetration depth rather than intrinsic tissue factors like hydration, control collections were performed on the underside of the skin without puncture, directly from the ISF-rich region of the dermis. Similar amounts of ISF were collected from both skin types, suggesting comparable hydration levels and hydraulic conductivities. Statistical analysis used a two-way ANOVA with post-hoc Tukey test. ns = not significant, ** = $p < 0.01$.

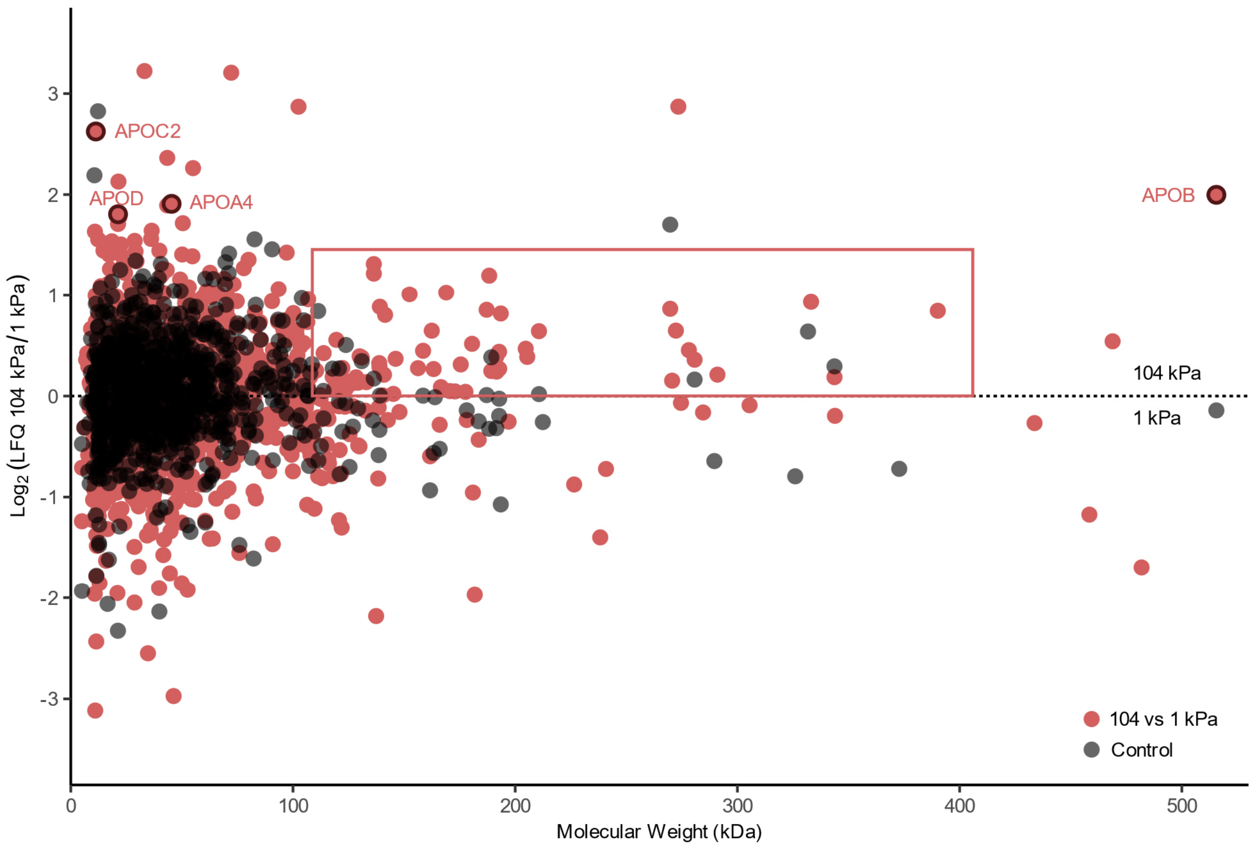


**Figure S9. Larger analytes are slightly over-represented in the ISF proteome when collected under high pressure.** The red box highlights the subtle tendency for high molecular weight proteins to be over-represented in the ISF proteome at 104 kPa compared to 1 kPa. Apoproteins, presumed to correlate with lipoprotein particle levels, were among the most over-represented at higher pressure. These trends, albeit subtle, favor a size exclusion model of analyte separation in the ECM over a filtration model. Controls were collected at 0 and 30 minutes without pressure, resulting in similar ISF except for a degree of evaporation at 30 minutes.

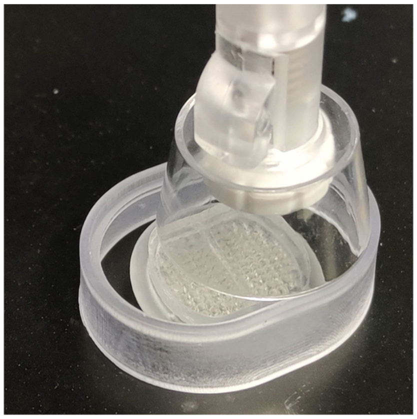

**Figure S10. Applicator stand for POP.** The stand allows the applicator to be deployed at a controlled height to strike the MAP. Additionally, the applicator can slide along the stand to strike the MAP at three different locations for more even penetrations.

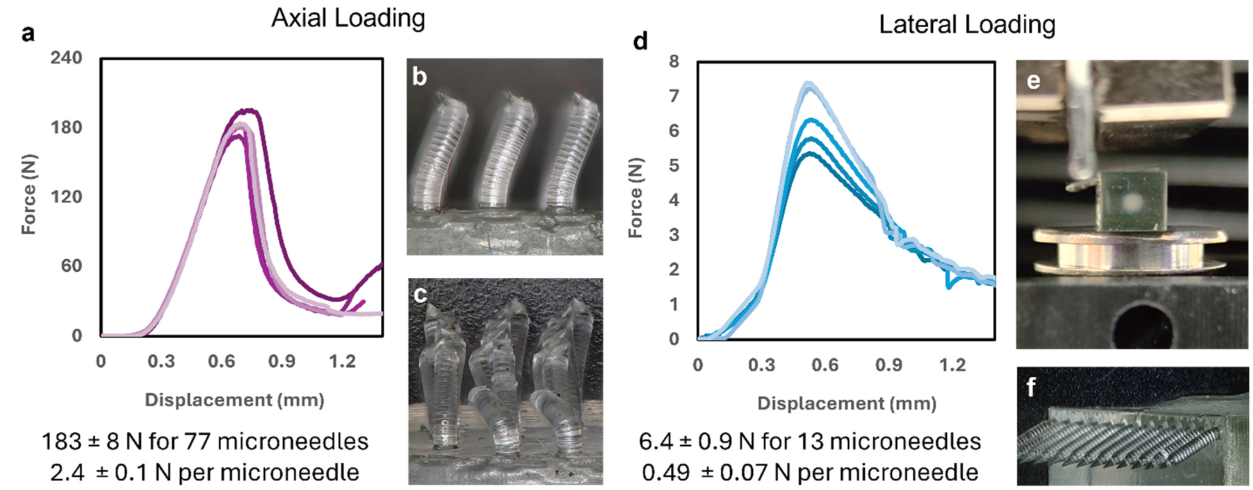


**Figure S11. Mechanical testing of solid MAPs. a** MAPs failed at 2.4 ± 0.1 N per microneedle under axial load (N=5). **b** Failure occurred through viscoelastic deformation at approximately 1.2 mm displacement. **c** With further displacement to approximately 1.7 mm, the microneedles continued to deform without fracture. **d** MAPs failed at 0.49 ± 0.07 N per microneedle under lateral load (N=5). **e** Lateral load testing applied force using the backside of a razor. **f** The failure mode was viscoelastic deformation without fracture, and the microneedles partially returned to upright position within minutes after testing.

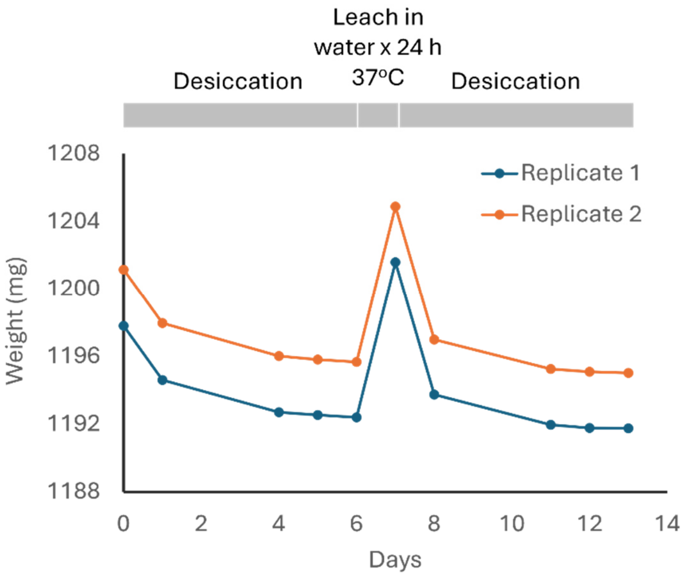


**Figure S12. Characterization of monomer leaching in POP.** The study followed ISO 20795-2. Leaching was performed in water at 37°C for 24 hours, shortened from the standard 7-day duration due to the device's 5-minute skin contact time. Version C (clinical) of the POP device was tested, comprising of a MAP mated to a collection plate. A water solubility of 0.598 ± 0.002 µg/mm³ was calculated from the dry weights pre- and post-leaching and the device volume estimated from its 3D model. The allowable limit per ISO 20795-2 is 5 µg/mm³.

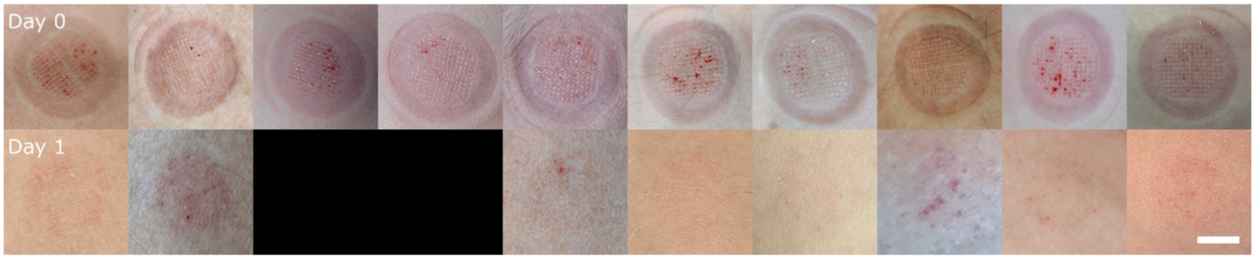


**Figure S13.** Photographs of POP puncture sites in human subjects immediately post-collection on day 0 (top row) and on day 1 (bottom row). Scale bar = 10 mm.

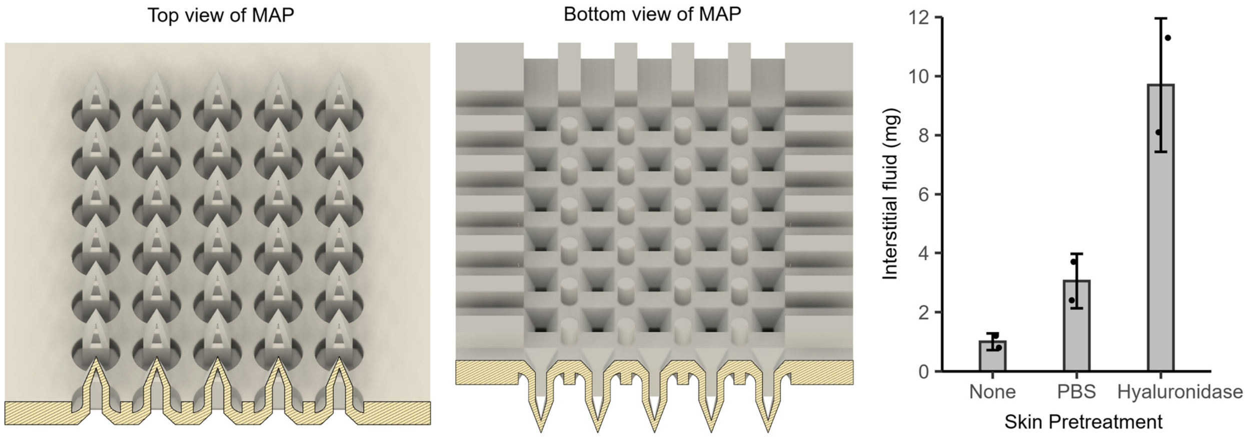


**Figure S14. Mimicking skin aging by pre-treatment with hyaluronidase.** To test the hypothesis that ISF flow increases with age due to ECM loss, intradermal injection was performed on *ex vivo* human skin with 10 μL of hyaluronidase (20-50 units), compared to sham injections and 10 μL of PBS. ISF collection from pre-treated skin demonstrated that degrading hyaluronic acid significantly increased ISF collection, similar to the trend observed between younger and older human subjects. The MAP used in this study had a monolithic design.

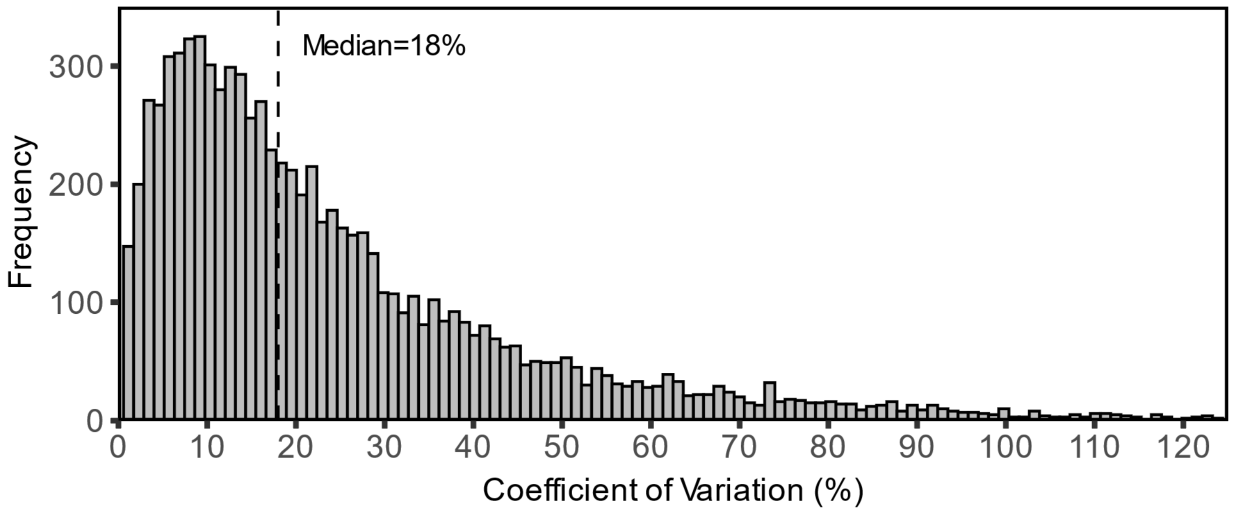


**Figure S15. Histogram of the coefficient of variation for 8335 triplicate label-free quantification measurements.** The median quantification error is 18%. In blood contamination analyses, a protein is considered contaminated if the error from contamination exceeds the measurement error. To establish an upper limit on errors from contamination, a stringent threshold of >10% was applied rather than >18%. Thus, the number of contaminated proteins estimated by the presented analyses is likely greater than would be observed in practice.

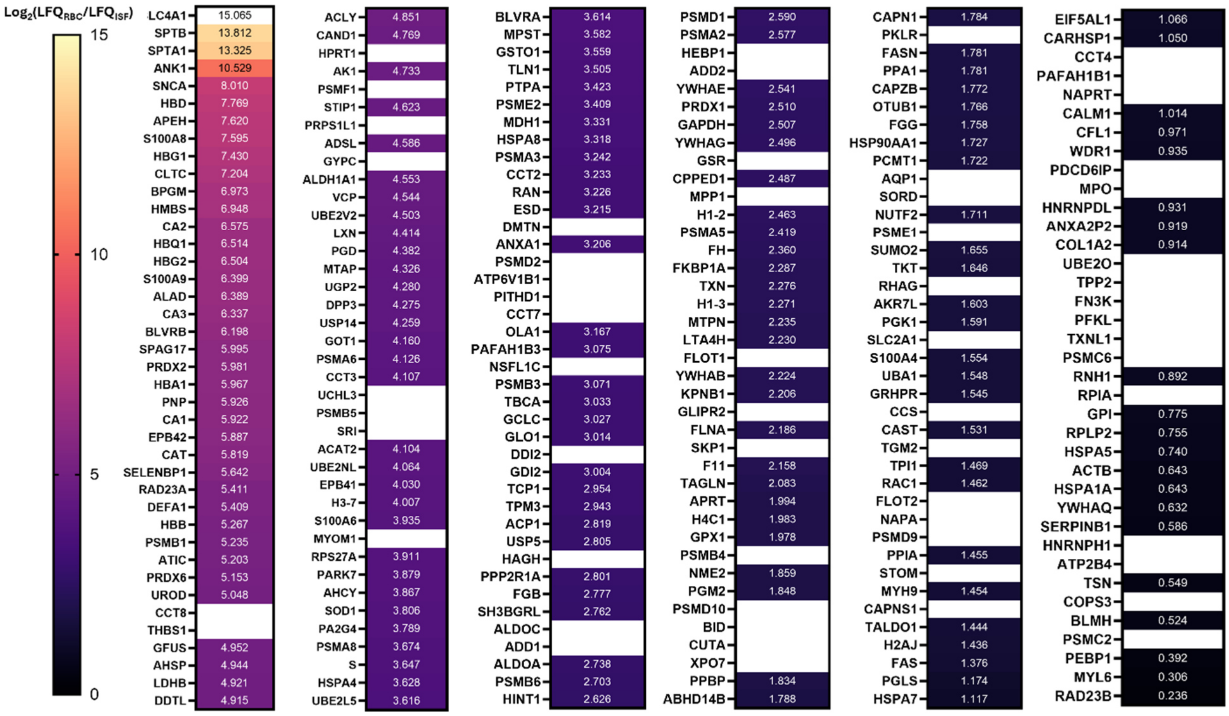


**Figure S16. A ranked list of RBC contaminant proteins, ordered from most to least contaminating by label-free relative quantification against ISF.** Proteins lacking LFQ ratios were only identified in RBC and were included because they may become measurable in ISF with increased blood contamination. To remove proteins in ISF with quantitation errors from RBC contamination, go through the list in order and remove each protein from ISF analysis until a pre-determined count has been reached.

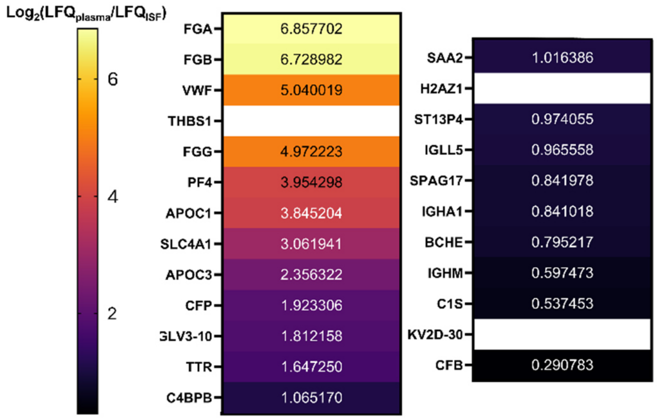


**Figure S17. A ranked list of plasma contaminant proteins, ordered from most to least contaminating by label-free relative quantification against ISF.** Proteins lacking LFQ ratios were only identified in plasma and were included because they may become measurable in ISF with increased blood contamination. To remove proteins in ISF with quantitation errors from plasma contamination, go through the list in order and remove each protein from ISF analysis until a pre-determined count has been reached.

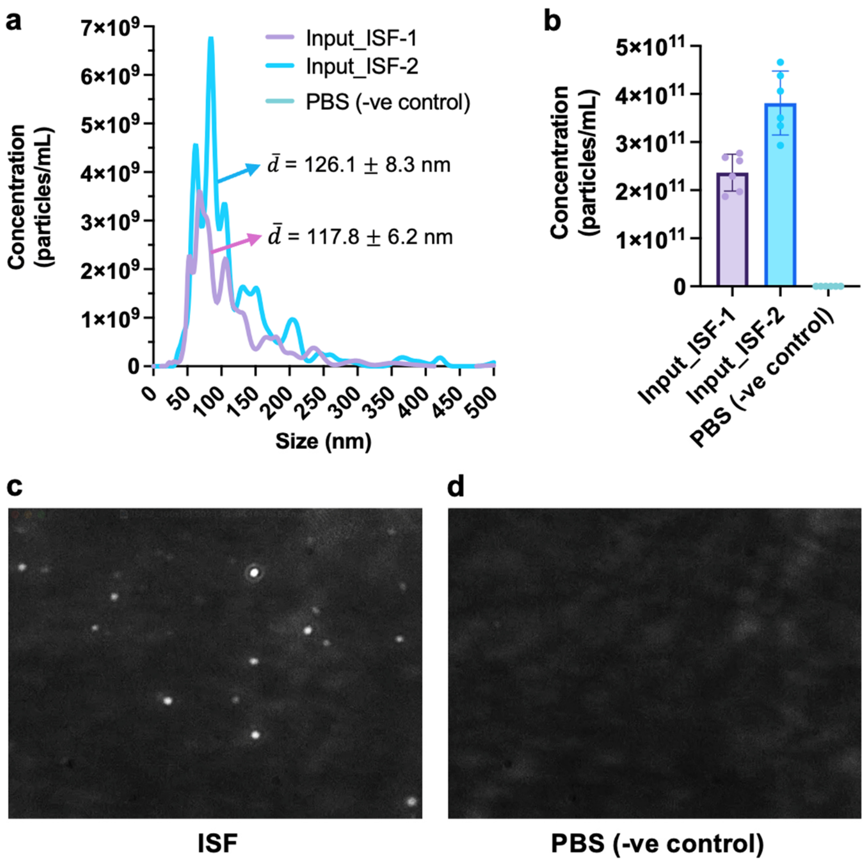


**Figure S18. Nanoparticle Tracking Analysis (NTA) of ISF from *ex vivo* Human Skin.** The presence of nanoparticles sized 50-250 nm in ISF suggests that apolipoproteins in ISF do not exist exclusively as the lipid-free form. In human plasma, up to 98% of the particles measured by NTA are lipoproteins, primarily VLDL and chylomicrons[1,2]. The assumption that apolipoprotein levels in ISF correlate with lipoprotein particle levels is plausible. **a** Extracellular vesicles and particles (EVPs) in ISF have a mean diameter of 110-130 nm. **b** EVP concentration in ISF ranges from 2.4 x $10^{11}$ to 3.8 x $10^{11}$ particles/mL, compared to 1-5 x $10^{12}$ particles/mL in plasma[1,2]. This is consistent with the lower representation of APOB and APOC in the ISF proteome relative to plasma in the mass spectrometry measurements. **c** Representative NTA micrograph of an ISF sample confirms the presence of nanoparticles. **d** Representative NTA micrograph of PBS, as negative control. Two biological replicates were performed.

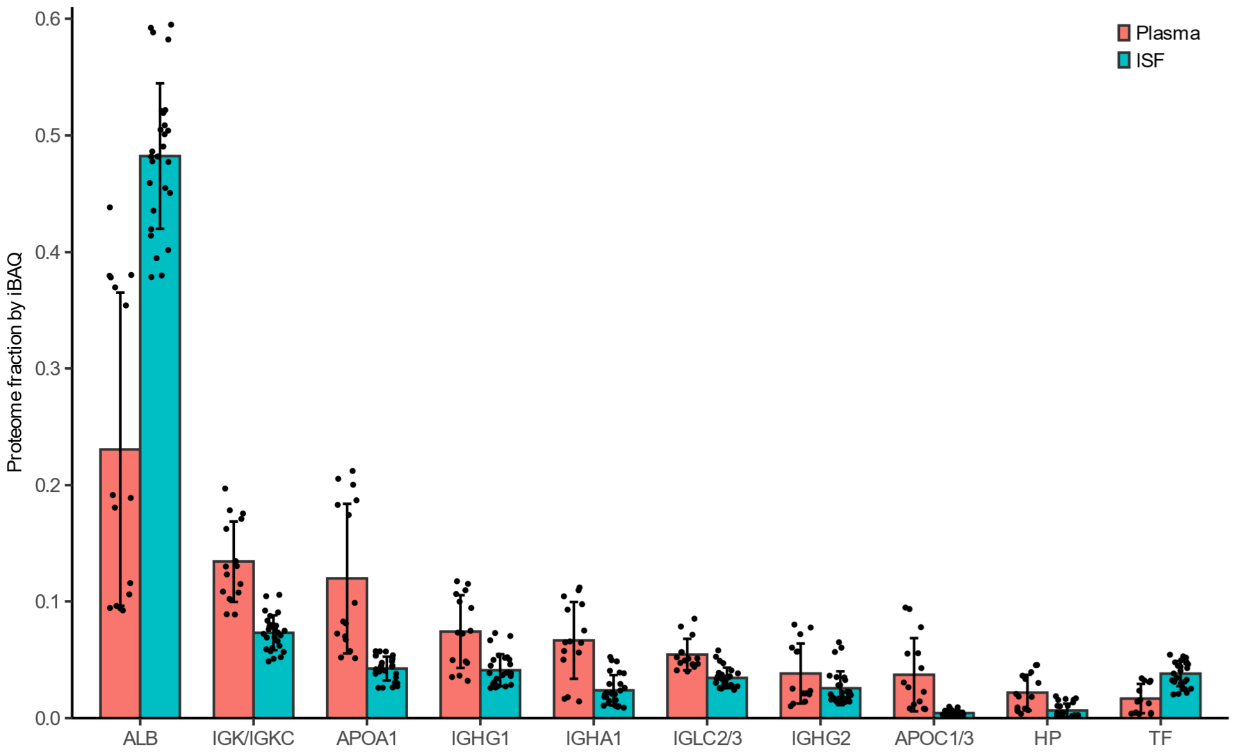


**Figure S19. Comparison of the most abundant proteins in human plasma and ISF.** Albumin (ALB) comprises a larger fraction of the ISF proteome than the plasma proteome, while apolipoproteins (APOA1, APOC1/3) make up a smaller fraction. The high variability in plasma among individuals reduces the accuracy in estimating the magnitude of the difference. For immunoglobulins, the difference between plasma and ISF is less clear, with considerable overlap in IgG1 (IGHG1), IgG2 (IGHG2), and lambda light chains (IGLC2/3).

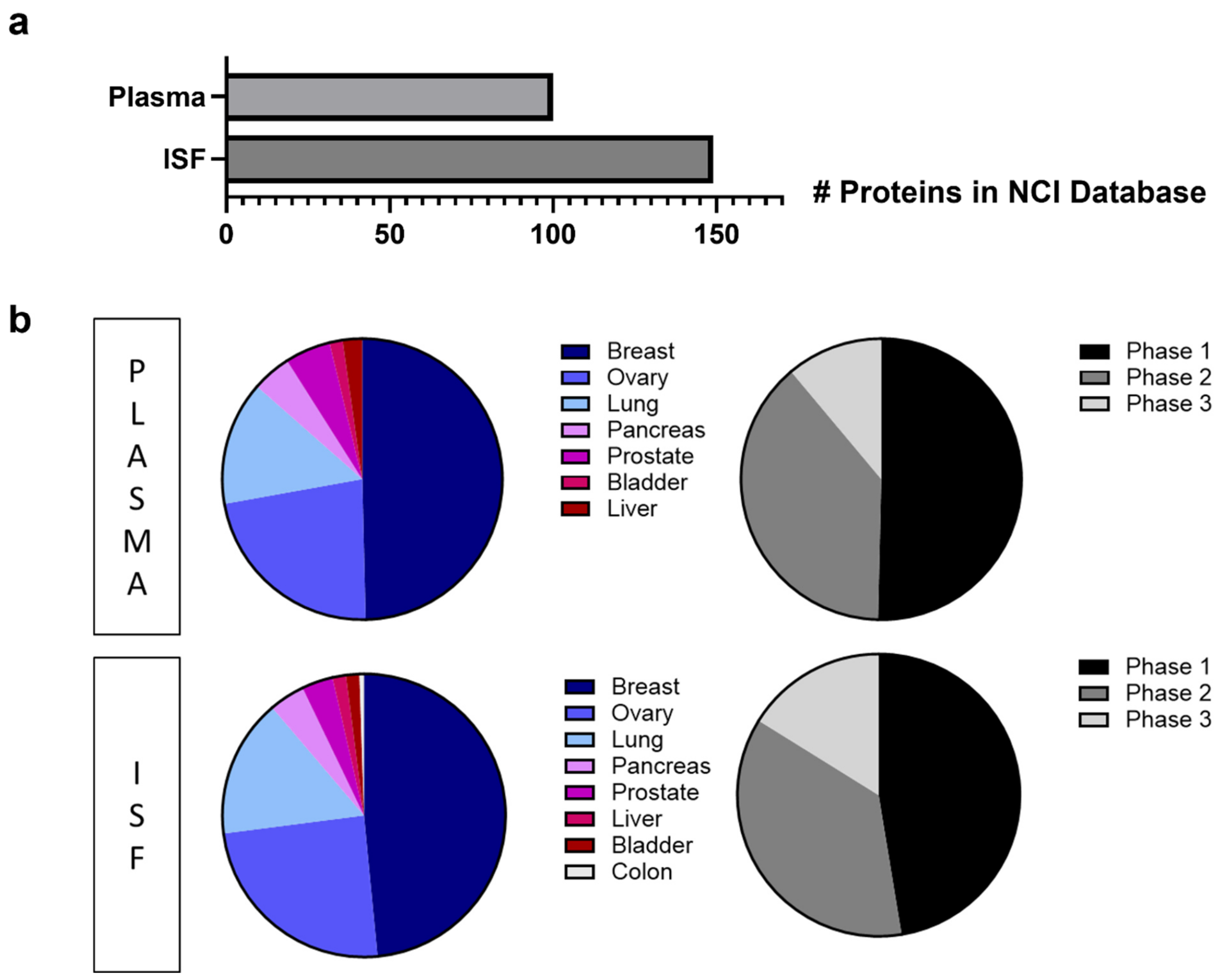


**Figure S20. Potential protein biomarkers in ISF. a** Comparison of proteins found in ISF and plasma that overlap with the NCI EDRN Biomarker database. **b** Distribution of protein biomarkers from the NCI database found in plasma and ISF. A majority of biomarkers are associated with breast cancer and are in phase 1 development.

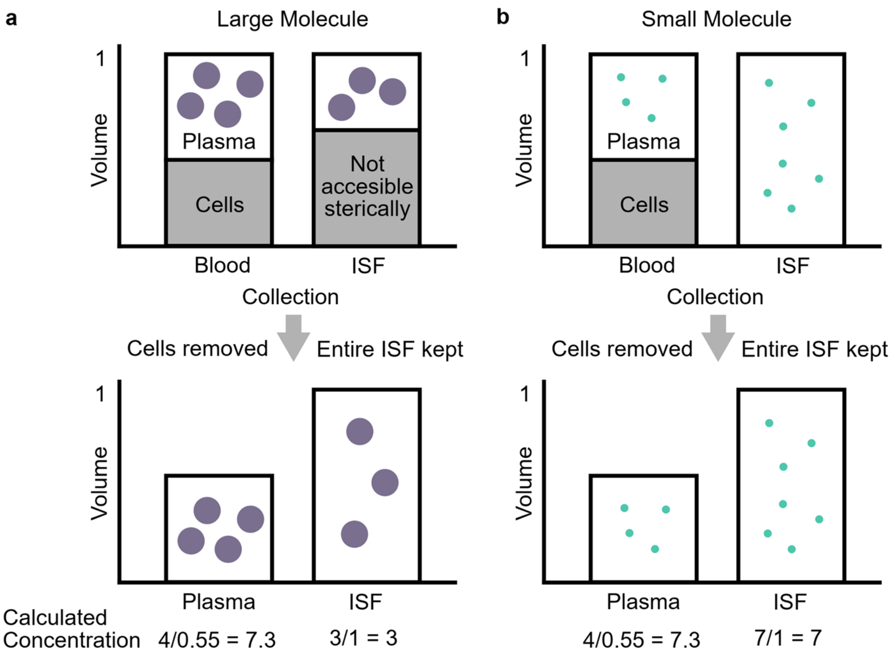


**Figure S21. Concentrations of large molecules in ISF are underestimated relative to plasma when the sterically inaccessible portion of ISF is not considered. a** When calculating analyte concentration in plasma, the inaccessible volume fraction in cells is routinely excluded. Analogously, there is an inaccessible volume fraction in ISF due to steric hindrance by the extracellular matrix (ECM), particularly for large molecules. However, this fraction has not been excluded in ISF calculations historically, leading to underestimation of concentrations relative to plasma. **b** In contrast, small molecules have little to no inaccessible volume fraction in ISF, allowing for accurate estimation of ISF analyte concentration relative to plasma.

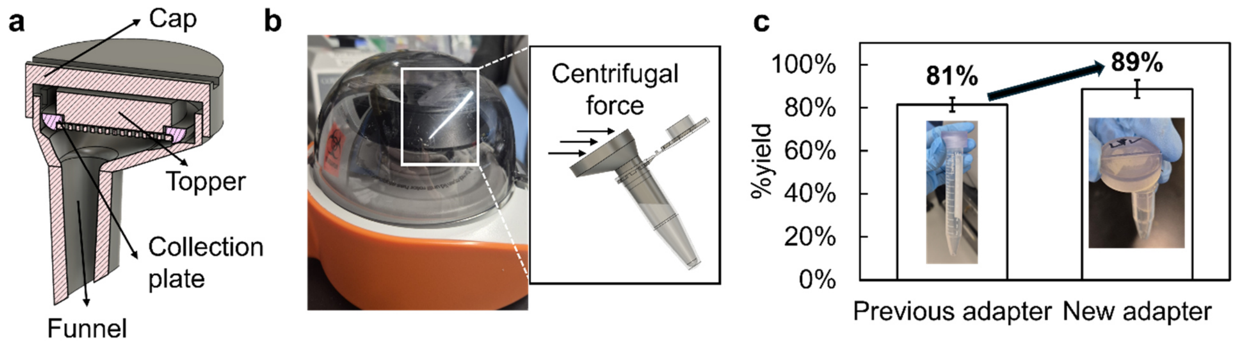


**Figure S22. Microcentrifuge adapter for ISF extraction from collection plate.** The adapter enhances POP's portability in clinical settings that lack full-sized centrifuges. **a** Adapter design features a tapered funnel for microcentrifuge tubes. **b** Its asymmetric design directs ISF into the funnel via centrifugal force. **c** The design achieves nearly 90% volume recovery from the plate.

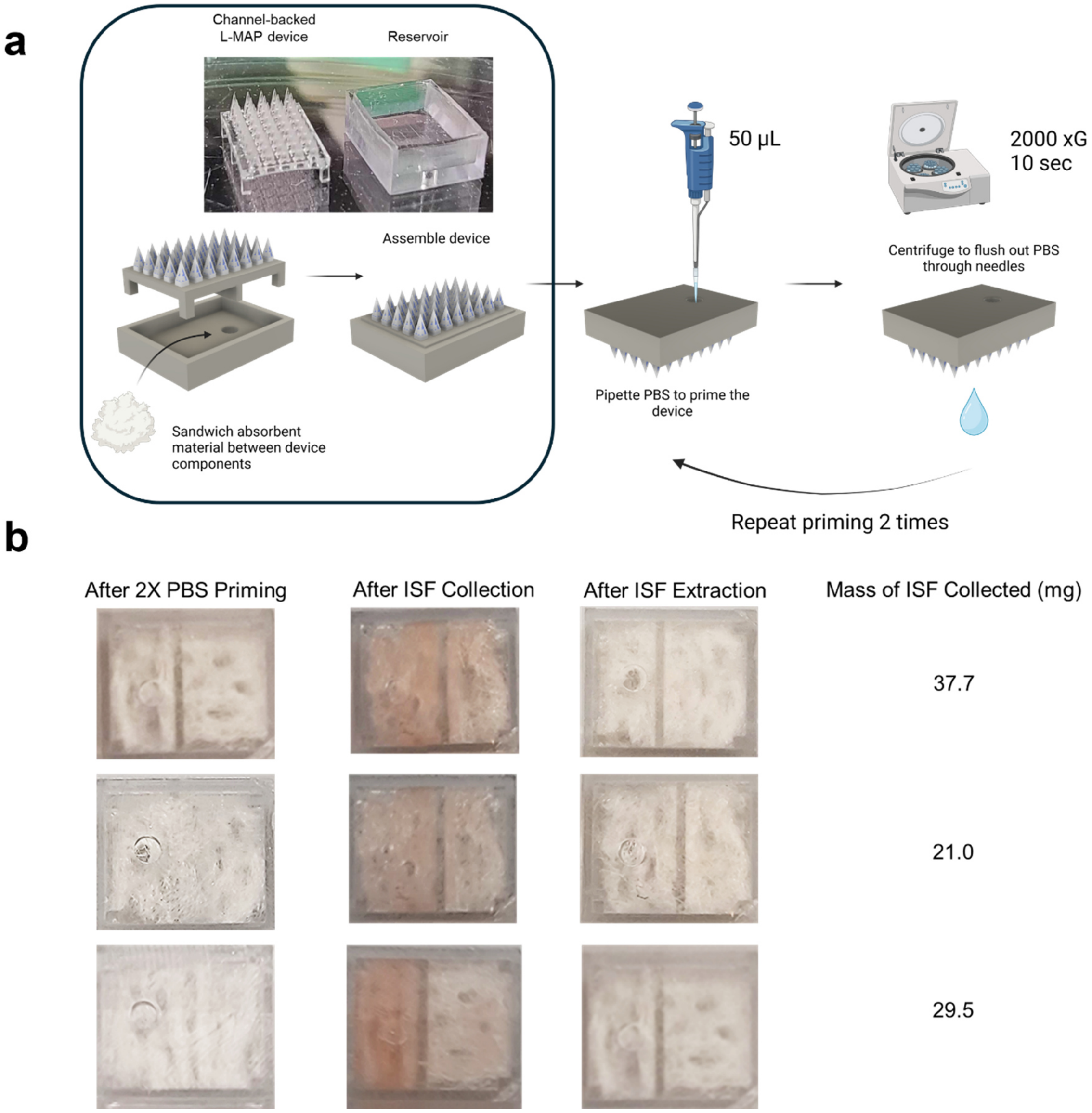


**Figure S23. Monolithic latticed MAP device. a** ISF collection process. An absorbent material such as gauze is sandwiched between the L-MAP and reservoir, which fit together. PBS primes the L-MAP, gauze retains the collected ISF, and centrifugation extracts ISF out of the gauze for analysis. **b** Micrographs of the L-MAP reservoir after PBS priming, ISF collection, and centrifugation. A spring-loaded applicator was used to puncture skin. ISF was collected over 5 minutes from porcine ear skin ex vivo, with pressure applied by a 500 g weight. 27.6 ± 8 mg of ISF was collected.

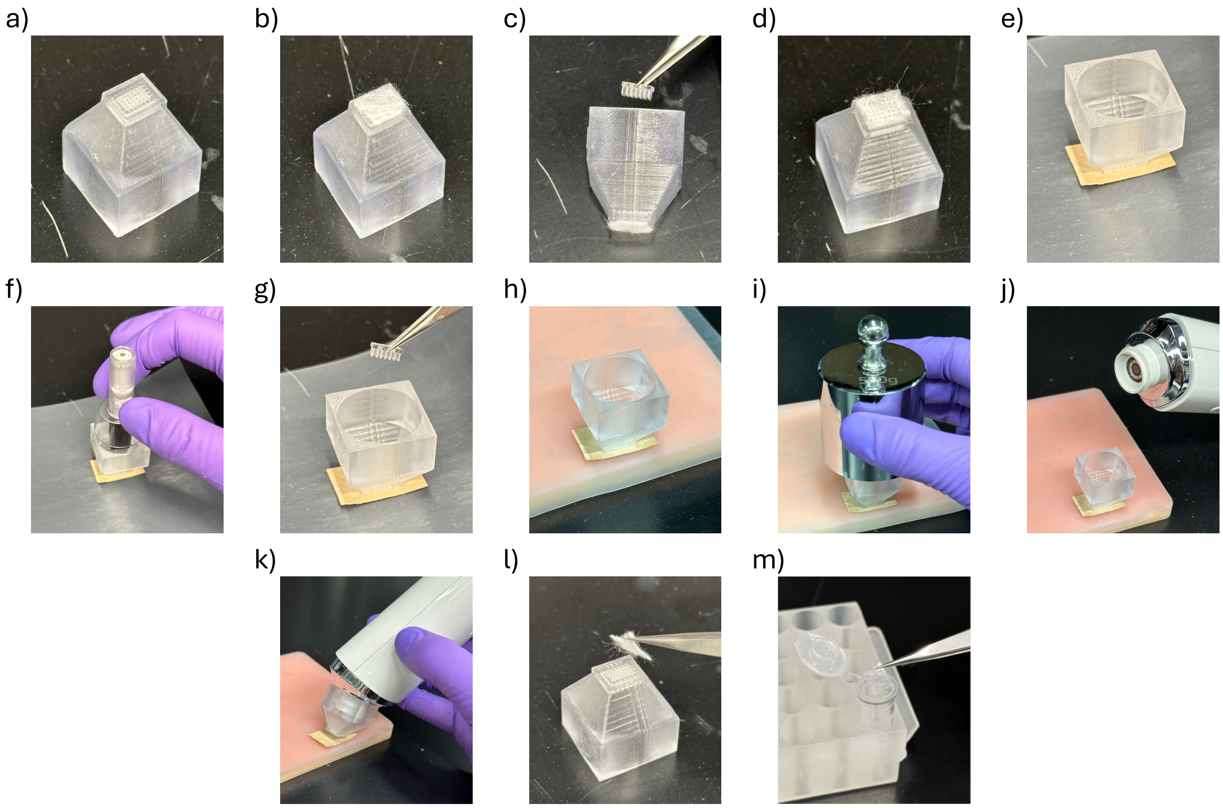


**Figure S24. ISF collection procedure in skin *ex vivo.*** a) ISF collection apparatus comprising of a collection plate affixed to a vacuum adapter. b) Gauze can optionally be adhered to the plate by friction to reduce operator variability in partial ISF loss when the collection apparatus is lifted from the skin. c) A MAP is placed into the collection apparatus and mated with the plate. d) If gauze is used, the gauze surrounding each exposed microneedle is pushed down using tweezers. e) The assembly is placed on excised skin on parafilm. The MAP retracts slightly into the plate upon skin contact. f) An applicator with a custom extender is placed on top of the collection apparatus and deployed three times to strike the MAP and puncture skin. g) The MAP is removed, ensuring that the collection apparatus remains aligned to the puncture sites. h) The setup including parafilm is transferred to a silicon pad to simulate subcutaneous fat. i) A weight is placed on the collection apparatus and balanced by hand to exert pressure. j) Optionally, a vacuum equipped with a 20x camera can be attached to visually monitor ISF retention during collection apparatus removal. k) The collection apparatus is slowly peeled away from the skin, and ISF is held in the plate perforations, or the gauze if used. The skin is typically held down with tweezers. l) If gauze is used, it is removed at this time and wiped against the plate to ensure complete collection of ISF. m) The gauze is placed into 100 µL of 200 mM ammonium bicarbonate in LC-MS grade water.

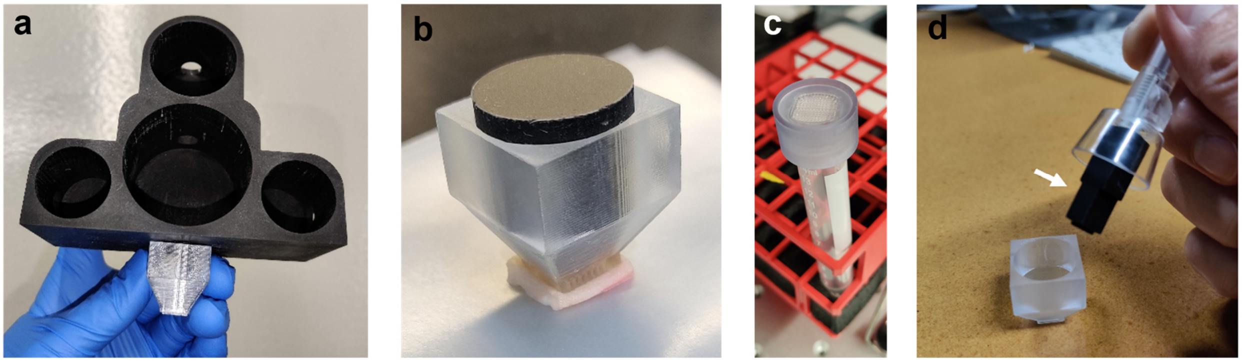


**Figure S25.** Experimental accessories. **a** Custom 3D-printed holder to allow for multiple calibrated weights to be fitted on top of the collection apparatus to apply pressure. **b** Custom 3D-printed insert that fits in the collection apparatus to minimize evaporation. **c** Custom adapter fitted on top of a 15 mL conical tube for centrifuging ISF out of a collection plate. **d** Custom extender on commercial applicator for striking a MAP from the top of the collection apparatus.

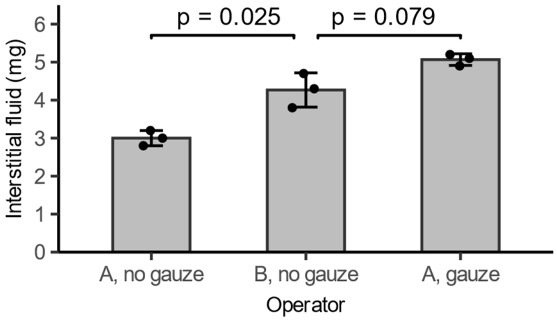


**Figure S26. ISF collection by POP varied by operator technique.** When removing the collection apparatus from the skin, ISF can be partially lost due to insufficient surface tension. Using gauze prevented ISF loss and reduced discrepancies between operators. ISF loss was 15-40% depending on operator. Version B of the POP device was used.

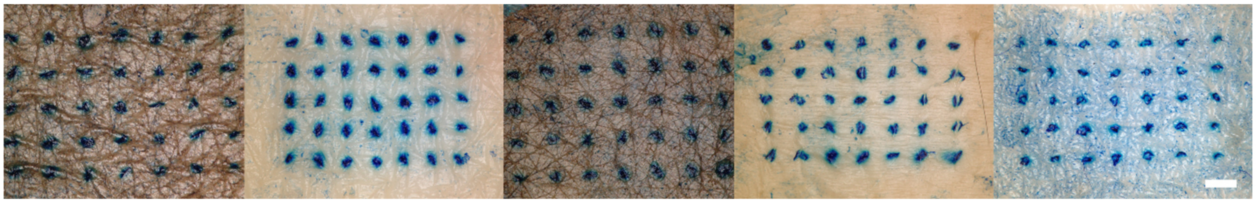

**Figure S27.** Examples of MAP puncture verification by methylene blue dye after *ex vivo* human skin experiments. Methylene blue stains the tissue where the stratum corneum has been breached. Scale bar = 1 mm.

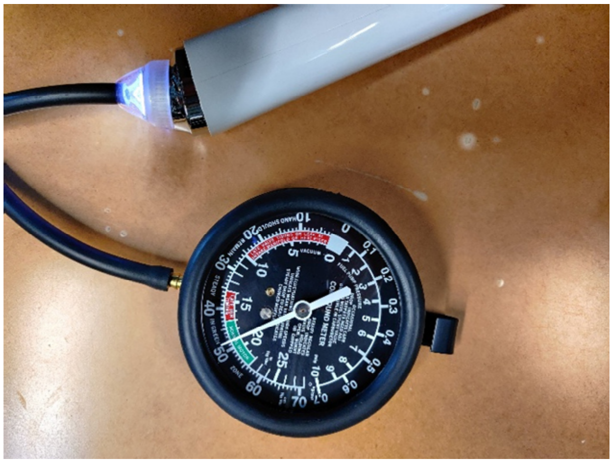

**Figure S28. Pressure measurement for commercial vacuum.** The commercial facial pore vacuum device applies -48 cm Hg, or -68 kPa, of vacuum on high setting.

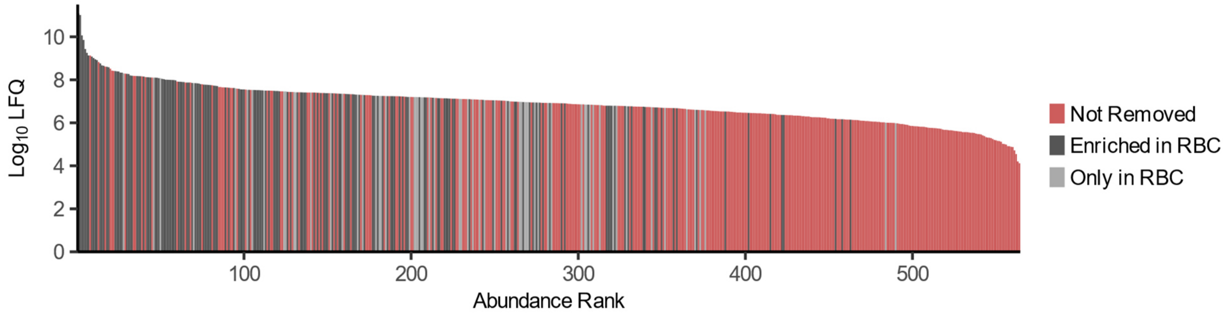


**Figure S29. RBC contaminant proteins visualized within the RBC proteome by abundance.** "Enriched in RBC" denotes RBC contaminant proteins found in both RBC and ISF that are more abundant in RBC. These proteins tend to be some of the most abundant proteins in the RBC proteome, as expected. "Only in RBC" denotes RBC contaminant proteins identified solely in the RBC proteome. These proteins tend to be in the middle rank of abundance among RBC proteins. These contaminants may appear in ISF as blood contamination increases beyond the level encountered in this study. Based on this data, the naïve strategy of removing the most abundant RBC proteins as contaminants would not achieve the same results as the strategy presented in this report. LFQ stands for label-free quantification.

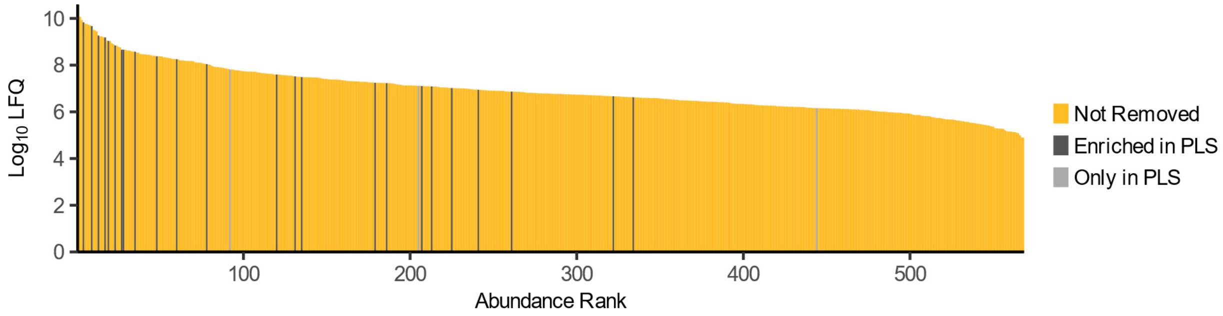


**Figure S30. Plasma contaminant proteins visualized within the plasma proteome by abundance.** "Enriched in PLS" denotes plasma contaminant proteins found in both plasma and ISF that are more abundant in plasma. These proteins tend to be some of the most abundant proteins in the plasma proteome, as expected. "Only in PLS" denotes plasma contaminant proteins identified solely in the plasma proteome. These contaminants may appear in ISF as blood contamination increases beyond the level encountered in this study. Based on this data, the naïve strategy of removing the most abundant plasma proteins as contaminants would not achieve the same results as the strategy presented in this report. LFQ stands for label-free quantification.

### Supplementary Tables

#### Table S1. Demographics in Human Study

| Characteristics | Participants |
|---|---|
| Age (years) | |
| 18 – 40 | 3 |
| 41 – 59 | 5 |
| 60+ | 2 |
| Sex | |
| Male | 6 |
| Female | 4 |
| Ethnicity | |
| Caucasian | 6 |
| Asian | 3 |
| Multiple | 1 |

#### Table S2. Negative control to rule out ISF contamination by skin surface proteins

| | Sham Collection | Human ISF Collection |
|---|---|---|
| ISF amount | - 0.6 ± 0.6 mg (N = 3) | 15.5 ± 7 mg (N = 10) |
| Raw protein concentration* | 29 ± 12 μg/mL (N = 3) | 3459 ± 1161 μg/mL (N = 9) |

*measured from ISF supernatant directly after centrifuging it from the collection plate. The sham collection was identical to the ISF collection, except without MAP skin puncture

#### Table S3. Comparison of immunoglobulin quantification with previous work

| | | Arevalo et al. (N = 3)[3] | | This work (N = 5) | |
|---|---|---|---|---|---|
| Immunoglobulin | Description | Accession | Plasma/ISF | Accession | Plasma/ISF |
| IgA1 | heavy constant alpha 1 | P01876 | 1.3 | P01876 | 1.8 |
| IgA2 | heavy alpha 2 | P0DOX2 | 1.0 | P0DOX2\|P01877 | 0.7 |
| IgD | heavy delta | P0DOX3 | 0.9 | P0DOX3\|P01880 | 1.2 |
| IgE | heavy epsilon | P0DOX4 | 1.3 | N/A | N/A |
| IgG1 | heavy gamma 1 | P0DOX5 | 0.9 | P0DOX5\|P01857 | 1.0 |
| IgG2 | heavy constant gamma 2 | P01859 | 0.8 | P01859 | 1.1 |
| IgG3 | heavy constant gamma 3 | P01860 | 0.9 | P01860 | 0.6 |
| IgG4 | heavy constant gamma 4 | A0A286YFJ8 (fragment) | 1.9 | P01861 | 0.3 |
| IgM | heavy mu | P01871 | 1.5 | P01871 | 1.5 |

Reference[3] employed TMT quantification. This work employed label-free quantification.

**Table S4. Demographics of human skin used for ex vivo experiments**

| Sample ID | Age | Gender | Site | Experiment |
|---|---|---|---|---|
| 1 | 61 | Male | Abdomen | Pressure Distribution |
| 2 | 61 | Male | Abdomen | Pressure Series |
| 3 | 67 | Male | Abdomen | Time Series |

**Table S5. Finite Element Analysis Skin Model Parameters**

| Parameter | Modeled value |
|---|---|
| Density | 1.02 g / cm[2,4] |
| Young's modulus* | 100 kPa[4–6] |
| Poisson ratio** | 0.48[7,8] |
| Yield strength*** | 15 MPa[9–11] |
| Ultimate tensile strength*** | 15 MPa[9–11] |
| Shear modulus | 5 kPa[12,13] |

*Literature values range 5 kPa – 20 MPa. A value in the middle of the range that results in simulated deformations similar to those observed experimentally was chosen.
**Based on agar phantom and pig skin estimates
***Estimates from human and rabbit skin range 3 – 27 MPa

**Supplemental Methods**

**Comparison of Solid and Hollow MAPs in POP ISF Collection**

Methods described in the ISF Collection section were followed using *ex vivo* human skin. The collection plate featured a 7 x 5 array of perforations, each 500 μm in diameter and 700 μm deep, without spacer pegs. The solid MAP consisted of a 7 x 5 array of cylindrical posts, 400 μm in diameter and 1200 μm long, topped with conical tips 1000 μm tall. The hollow MAP featured a 7 x 5 array of microneedles, 400 μm in diameter with 200 μm wide channels and 2200 μm in length. All components were made from Keysplint Hard. Vacuum was applied during device removal to minimize ISF loss.

***Hyaluronidase Skin Pretreatment***

The MAP featured a monolithic design with hollow microneedles. Each microneedle had a 440 x 400 x 500 μm (L x W x H) rectangular body with 200 μm wide channels and a 1000 μm tall tip with two side openings. Each channel connected to a reservoir in the backing for ISF collection. On the skin side, each microneedle was seated in a circular inset of 800 μm diameter, mimicking the design of the POP collection plate. The MAP was affixed to a vacuum adapter to form a single unit, and methods described in the ISF Collection section were followed using *ex vivo* human skin. Removal of the MAP before pressure application was omitted due to the monolithic design. Vacuum was applied during device removal to minimize ISF loss. Prior to ISF collection, a 31G needle (UltiCare, Insulin Syringe, SY5751509436, Amazon) was used to inject 2 μL of PBS or hyaluronidase at five sites for a total of 10 μL in the skin. Hyaluronidase (from bovine testes, 400-1000 u/mg powder, H3506-100MG, Sigma Aldrich) was prepared at 5 mg/mL in PBS. Sham injections were performed for the control. The skin was massaged gently to distribute the injected payload before ISF collection.

**Nanoparticle Tracking Analysis**

Methods described in the ISF Collection section were followed using *ex vivo* human skin to obtain ISF. Version B of the POP device was used. Nanoparticle tracking analysis (NTA) was performed using a NanoSight NS300 with software build 3.44 (Malvern Panalytical) to determine the concentration and size distribution of nanoparticles in ISF. Each ISF sample was first diluted in 100 μL of PBS directly on the collection plate and centrifuged into a tube. 5 μL of the diluted sample was further diluted in 495 μL of PBS for NTA measurements, with a total dilution factor of approximately 1000x. The diluted sample was loaded into a 1-mL syringe for analysis. The NanoSight instrument was maintained at 25°C, and each measurement sequence comprised three runs, each capturing a 30-second video. A syringe pump was used at a speed of 20 units, with the camera level set to 14 and the detection threshold at level 5. This measurement sequence was repeated twice for each sample. Reported statistics represent the average of six runs. Between measurements, the infusion lines and microfluidic chamber were washed with PBS to prevent cross-contamination.

**Mechanical Testing**

Mechanical evaluation of MAPs was conducted using an MTS Criterion Electromechanical Test System with either a 1000 N or a 100 N load cell depending on the force requirement. MAPs were adhered to a scanning electron microscope pin stub mount (#16111, Ted Pella Inc.) by superglue and placed on a custom holder. For axial loading, the MAPs were mounted upright, and a metallic plate compressed the tips at 50 μm/s. For lateral loading, a single row of microneedles was 3D-printed and mounted sideways. The custom holder was taped to the testing stage to prevent lateral movement. A grip attachment with a clamped razor was used to apply blunt force to the side of the microneedles at 50 μm/s via the backside of the razor. Micrographs were taken using an Olympus DSX1000 microscope to show the failure mode.
